# AUF-1 knock down in mice overarches butyrate driven hypo-cholesteraemia by conjuring AUF-1-Dicer-1-miR122 hierarchy

**DOI:** 10.1101/2022.06.13.496022

**Authors:** Oishika Das, Jayanta Kundu, Atanu Ghosh, Anupam Gautam, Souradeepa Ghosh, Mainak Chakraborty, Aaheli Masid, Samiran Sona Gauri, Debmalya Mitra, Moumita Dutta, Budhaditya Mukherjee, Surajit Sinha, Moumita Bhaumik

## Abstract

This discourse probes the mechanistic molecular details of butyrate action in maintaining host-cholesterol balance. Hepatic miR122 being the most indispensable regulator of cholesterol metabolic enzymes, we studied upstream players of miR122 biogenesis in the presence and absence of butyrate in Huh7 cells and mice model. We showed that butyrate treatment caused upregulation of RNA-binding protein, AUF-1 resulting in RNase-III nuclease, Dicer-1 instability, and significant diminution of miR122. We proved its importance of AUF-1 and sequential downstream players in AUF-1-knock-down mice. We synthesized unique self-transfecting GMO (guanidinium-morpholino-oligonucleotides) linked PMO (Phosphorodiamidate-Morpholino Oligonucleotides)-based antisense reagent and injection of which in mouse caused near absence of AUF-1 coupled with increased Dicer-1 and miR122, and reduced serum cholesterol regardless of butyrate treatment indicating that butyrate acts though AUF-1. The roster of intracellular players was as follows: AUF-1-Dicer-1-miR122 for triggering butyrate driven hypocholesterlaemia. To our knowledge this is the first report linking AUF-1 with cholesterol biogenesis.

## Introduction

The gut microbiome is endowed with the special ability to govern host metabolic function which cannot be accomplished by the host itself. Gut microbiota on fermentation of non-digestible carbohydrates produce short-chain fatty acids (SCFAs) like acetate, propionate, and butyrate at a ratio of 3:1:1 ^1^. There is colossal evidence that gut-derived SCFAs display overarching effects on health and diseases^2^. The benefits of butyrate are particularly well-researched as a therapeutic agent against inflammatory diseases, neurological disorders and others ^3^. Butyrate is also reported to mediate the host cellular energy metabolic landscape ^4^. Studies in mouse models have shown that a diet supplemented with butyrate prevents high fat diet (HFD) induced obesity, by down-regulating the expression and activity of PPAR-γ ^5^, while the intake of dietary polysaccharide pectin that produces butyrate after fermentation in gut, inhibits cholesterol uptake and promotes cholesterol efflux from enterocytes in Apo E^−/−^ mice ^6^. There are different types of lipoproteins (HDL, LDL, VLDL, chylomicron) consisting of different percentage of cholesterol, protein, triglyceride and phospholipids. Cholesterol orchestrates a number of important cellular functions like membrane fluidity, steroid hormone synthesis, bile acid synthesis, while its intermediate metabolic precursor, 7-dehydrocholesterol serves as a precursor of vitamin D ^7^. Dysregulation of cholesterol metabolism is associated with cardiovascular disease, neurodegenerative diseases, diabetes and cancer ^8–10^. Cholesterol homeostasis at the cellular level is governed at multiple layers like biosynthesis, catabolism and efflux, the latter pathway is accomplished by ATP-binding cassette transporter A1 (ABCA1) ^11^. These distinctly-different, linked paradigms are converged to a singular framework of host cholesterol homeostasis under the influence of butyrate.

At the cellular level, the regulation of cholesterol biogenesis and efflux is governed by microRNAs, the important ones include miR122 and miR27 respectively ^12 13^. MiR-122, making up 70% of the total hepatic miRNA ^14^ contributes to the maintenance of the adult liver phenotype ^15^. Mice treated with antagomir-122 shown to modulate cholesterol metabolic genes ^12^. Additionally, miR122 regulates tight junction protein, occludin ^16^, important in maintaining mucosal barrier and junctional integrity of hepatic tissue ^17^. Like most miRNAs, miR122 are generated from the primary precursor, pri-miR122 by nuclear RNaseIII Drosha; the resultant pre-miR122 undergoes processing with the help of Dicer1, another RNase III nuclease that processes the pre-miRNA into a 22-bp double-stranded RNA ^18^. Dicer1 mediated miR122 maturation is a critical step in miR122 biogenesis which occurs in the cytoplasm.

In mammals, the Dicer1 mRNA is destabilized by the presence of AU-rich elements (AREs) in their 3’-untranslated regions ^19^. The protein, ARE/poly(U)-binding/degradation factor-1 (AUF1) binds to many ARE-mRNAs and assembles other factors necessary to recruit the mRNA degradation machinery ^20^ which is a family of RNA binding proteins with four spliced variants: AUF1^p45^, AUF1^p42^, AUF1^p40^ and AUF1^p3719^. The AUF1 interacts with the 3ʹUTR of Dicer1 mRNA and causes mRNA destabilization reducing its half-life ^19^. AUF1 KO mice develop chronic dermatitis ^21^ and are prone to septicaemic shock ^22^. Importantly, AUF1 is found to decay mRNA of sphingosine kinase1 ^23^ which produces Spingosine-1-phospahte (S1P), a biomarker of sepsis ^24^ and other inflammatory diseases ^25^. Absence of AUF1 thus increases S1P which causes rapid expansion of inflammatory signals ^26^.

This study is designed for the conflation of gut derived butyrate flux on metabolic phenotype encompassing miRNAs, Dicer1, AUF-1, cholesterol metabolic enzymes and the ultimate testimony of cholesterol homeostasis. We explored domains like expression of cholesterol synthesis, catabolic genes and efflux protein in the presence and absence of butyrate. We showed that butyrate modulates selective isoforms of RNA binding protein AUF1 leading to down-regulation of Dicer1 and miR122, resulting in decrease in cellular cholesterol. To provide further leverage in our understanding, we had synthesized an unique self-transfecting GMO (guanidinium morpholino oligonucleotides) linked PMO (Phosphorodiamidate Morpholino Oligonucleotides)-based antisense reagent, to knock down AUF-1 in mice. Using AUF-1 knock down mice we provided crucial evidence that butyrate indeed exploits AUF1 to accomplish its cellular effect. The above discourse led to germane a new element of our understanding on the role of AUF1 in cholesterol metabolism. Narratives from our experimental studies when considered in tandem, culminated in the following trajectory of butyrate action “butyrate-AUF1-Dicer1-miR122-cholesterol metabolic enzymes-cholesterol” which is tentatively posited as “axis”.

## Materials and Methods

### Propagation of Huh7 cells, estimation of cellular protein and cholesterol

Huh7 cells were cultured in DMEM supplemented with 10% FCS and 50 µg/ml gentamycin at 37° C with 5% CO_2_. Cellular viability was assayed by MTT. Cholesterol estimation was performed using Amplex Red Cholesterol Assay kit. An aliquot of cell lysate was used for protein estimation using Pierce^TM^ BCA Protein Assay Kit following manufacturer’s protocol.

### Mice and Animal Ethics

Pathogen free C57BL/6 mice (6 weeks old) were procured from institutional animal facility of ICMR-NICED, Kolkata, India. All the protocol for the study was approved by the Institutional Animal Ethics Committee of ICMR-NICED, Kolkata, India, (NICED/BS/MB-001/2019). All mice were housed under a 12-hour light/dark cycle at controlled temperature. Experiments have been carried out in accordance with the guidelines laid down by the committee for the purpose of control and supervision of experiments on animals (CPCSEA), Ministry of Environment and Forests, Government of India, New Delhi, India.

### Dietary supplementation of sodium butyrate

The dietary supplementation studies were performed as reported earlier with minor modification ^27^. Briefly, a group of 10 adult C57BL/6 mice (divided as 5 in each group) were fed with HFD having 20% kcal protein, 20% kcal carbohydrate, 60% kcal fat and trace amount of cholesterol (0.007% w/w). After 4 weeks of HFD feed, 5 animals from the group were selected randomly and was further fed with 5% sodium butyrate (w/w) (Sodium butyrate in solid form were thoroughly blended into HFD) for 15 days (HFD-butyrate-mice). The remaining mice were continued with HFD for 15 days (HFD-mice). Fresh diet was replenished in alternate days. Age matched mice fed with regular chow diet served as normal group (chow-mice).

### Antibiotic treatment

The endogenous intestinal microbiota of 4-week old chow fed mice was depleted by gavage with broad spectrum antibiotics over 7 days ^28^. The antibiotics solution consisted of ampicillin, metronidazole and vancomycin diluted in sterile water. Mice received 200 mg/kg of ampicillin and metronidazole and 100 mg/kg of vancomycin once a day. Henceforth, the antibiotic treated mice are called Abx-mice.

### Probiotic treatment

Probiotic (each capsule containing 50 million Lactic acid bacteria, 2 million *Clostridium butyricum*, 30 million *Steptococcus faecalis* and 1 million *Bacillus mesentericus*) treatment was done as described previously with minor modifications ^29^. Briefly, a group of 20 mice (4 mice in each cage) were fed with cocktail antibiotic for 7 days and 5 mice were subjected to bowel cleansing with 1.2 ml of poly ethylene glycol (PEG) solution for each mouse prior to probiotic administration. Since mice are coprophagous, they were placed in clean cages having wired net at the bottom of the cage to prevent feeding of faeces. Faeces and blood (from tail vein) was collected before bowel cleansing. PEG solution contained PEG 3350 (77.5 g/L), sodium chloride (1.9 g/L), sodium sulfate (7.4 g/L), potassium chloride (0.98 g/L) and sodium bicarbonate (2.2 g/L) diluted in sterile tap water and was divided in five equal doses (200 µl/ mice/ dose) that were administered by oral gavage at 30 min intervals after a 2 h fasting (free access to water). The PEG solution and the inoculum were provided 24 h after the last antibiotic gavage. Each capsule of probiotic was suspended in 1 ml water and two inoculations of probiotics (200 μl) were administered *via* oral gavage to Abx-mice, on every alternate day till 2 weeks. The mice were transferred to sterile cage and straw. All animals received autoclaved deionised water and chow diet ad libitum for next 14 days. A group of 5 Abx-mice were continued with antibiotic feeding along with 5% sodium butyrate supplemented with chow diet for next 14 days (Abx-butyrate-mice). Another group of 5 Abx-mice were continued with antibiotic treatment for next 14 days. An age matched mice (5 in number) fed with normal chow diet without antibiotic treatment throughout the period of 21 days served as a control (chow-mice).

### MiR122 overexpression in mice

MiR-122 was overexpressed in mouse liver as described earlier ^30, 31^. The miR-122 expression plasmid or empty plasmid (mock) was injected through the tail vein of butyrate treated mice at a dose of 25 µg DNA in 100 µl saline/ mouse. Mice were sacrificed on day 4 post injection and serum cholesterol and miR122 expression in liver was measured.

### Mice faecal samples collection

Fresh faecal samples of all mice were collected between15:00~17:00 p.m. to minimize possible circadian effects. Samples were collected into empty sterile microtubes on ice and stored at −80 °C within 1h for future use.

### Faecal DNA extraction and determination of butyryl-coenzyme A (CoA):acetate CoA-transferase (ButCoAT) gene abundance by qPCR

The QIAamp DNA stool minikit (Qiagen), was used to extract DNA in QIAcube (Qiagen) from 40 mg of faecal pellet from each mouse. The DNA was measured in QIAxpert spectrophotometer (Qiagen). To compare the relative abundance of bacteria having *butCoAT* gene in faecal DNA of mice, qPCR analysis was carried out and quantified by SYBR green and normalized the data to total bacterial abundance using 16S rRNA-bacterial primers ^32, 33^. The sequence of the primers used is listed in Table S1.

### Measurement of butyrate in faeces by LC-MS

Faecal content of each mice (50 mg wet weight) was dissolved in distilled water at 1:10 (w/v) and homogenized in a Dounce homogenizer. Then, the content was centrifuged at 10,000g for 10 m and the supernatant was filtered through 0.45µm syringe filter. The supernatant was further diluted in LC-MS grade distilled water upto a final volume of 1 ml and were subjected to LC-MS analysis for SCFA as described by Cheng et al ^34^. In brief, a calibration curve was generated by using varying calibration of internal standards of butyrate from 2µmoles to 50µmoles. The LCMS separation was performed on the Agilent 1290 Infinity LC system which was coupled to Agilent 6545 Accurate-Mass Quadrupole Time-of-Flight (QTOF) with Agilent Jet Stream Thermal Gradient Technology with electrospray ionization (ESI) source. The suitable MS parameters were optimised and the high resolution mass spectra were obtained by performing the analysis in negative ionisation mode. The Chromatographic separation was achieved on Agilent ZORBAX SB-C18 column (2.1 × 100 mm, 1.8 µm) as stationary phase. The mobile phase consisted of a linear gradient of 0.1% (v/v) aqueous formic acid (A) and Methanol (B): 0–1.0 min, 0 % B (v/v); 1.0–3.0 min, 0–30% B (v/v); 3.0–6.0 min, 30-100% B (v/v); 6.0–12.0 min, 100% B (v/v); 12.50–15.0 min, 0% B. The column was reconditioned for 5 min before next injection. 0.2 mL/min flow rate and a varying injection volume was used for analysis. The UPLC system assembled with a Diode array detector (DAD) and an auto sampler. The peak area of each SCFA was used to calculate the amount of SCFA present which was further normalized by the injection volume. The data was represented as the amount of butyrate present per gram of faeces.

### Preparation of liposome and loading of cholesterol to Huh7 cells

Liposomes was prepared using 22-NBD-Cholesterol [22-(N-(7-Nitrobenz-2-Oxa-1,3-Diazol-4-yl)Amino)-23,24-Bisnor-5-Cholen-3β-Ol] as described ^35^. Briefly, 22-NBD-Cholesterol (1mg), Cholesterol (4.8mg) and Phosphatidylcholine (8mg), were dissolved in 1ml of chloroform. The solvent was evaporated to obtain a homogeneous thin lipid film and put into a 100 ml round-bottom flask, followed by solvent removal by rotary evaporation under reduced pressure. The lipid film was then hydrated in serum free DMEM and sonicated (Microson Ultrasonic cell disruptor with a Misonix 2-mm probe) at 4°C three times for 1 min each at maximum output and filtered with 0.2 µm Millipore filter ^36^. Huh7 cells were plated in 96-well plate at a cell density of 10^5^cells/well. Cells were equilibrated with 22-NBD-cholesterol-liposome for overnight. After incubation with liposome, cells were washed with 1XPBS and were subjected to treatment with or without butyrate.

### Cholesterol efflux assay

The cholesterol efflux assay was performed as described ^37^. Briefly, cholesterol loaded cells were washed with 1XPBS and then treated with butyrate for next 24 h. Following treatment with or without butyrate, cells were washed with 1XPBS and incubated in serum free DMEM for next 18 h. After specified time of incubation, cells were treated with 1 µg/ml, 5 µg/ml and 20µg/ml HDL respectively for 4 h to induce efflux. Thereafter, cell supernatant was collected and cells were lysed using methanol. The fluorescence intensity (FI) of 22-NBD-cholesterol in the medium and cell lysate was detected by MT-600F fluorescence microplate reader (Corona Electric, Hitachinaka, Japan) using 469 nm excitation and 537 nm emission filters in a black polystyrene 96-well plate (Costar, Corning Incorporated, USA). The percent cholesterol efflux was calculated as follows: % cholesterol efflux= (FI medium X100)/ (FI medium + FI lysate).

### GMO-PMO treatment to knock down AUF1

The GMO-PMO was dissolved in saline to make a concentration of 1 mM which was equivalent to 8.2 mg/ml. In vivo knock down of AUF1 was carried out in mice by injecting AUF1 GMO-PMO (AUF1-MO) through tail vein on day 1 and day 7 at a dose of 3 mg/kg body weight. The effective dose was previously determined based on AUF1 expression in liver by a pilot study. Another group of AUF1-MO injected mice were fed with 5% butyrate supplemented chow diet from day 1 and continued till day 14. Blood was collected from tail vein on day 7 and day 14 and for cholesterol estimation. Mock GMO-PMO treated animals are called as scramble-MO controls. Another group of scramble-MO were fed with butyrate for 14 days post first dose injection. Animals were sacrificed after day 14 and tissue were collected for further analysis.

### Statistical Analysis

All values in the figures and text are expressed as arithmetic mean ± SEM. Data were analyzed with GraphPad Prism Version 8.01 software and statistical significance between groups was determined using unpaired student’s t-test. The *p* values of *<*0.05 were considered statistically significant. In the experiment involving Western blot, the figures shown are representative of at least 3 experiments performed on different days.

## Results

### Butyrate is more effective than propionate and acetate in reducing cellular cholesterol and inhibits HMGCR

To witness which one of the SCFAs may interfere in cholesterol biogenesis with fidelity, we undertook an initial approach in the cell line Huh7, although most of our obvious thrust is in the mouse model. The Huh7 cell-line is considered essentially as a model for studying hepatic processes and therefore, it was used for this investigation. To assess the effects of SCFAs on cellular cholesterol biogenesis, Huh7 was treated with increasing concentration (0-20mM) of either sodium butyrate (butyrate) or sodium propionate (propionate) or sodium acetate (aceatate) for 24h following which cellular cholesterol was measured. The concentration of SCFAs used in the study based on the earlier reports ^38^. It was observed that unlike propionate and acetate (Fig S1), 5mM, 10mM and 20mM butyrate treatment caused 25%, 40% and 75% decrease in cellular cholesterol, respectively (Fig S1A inset). Since butyrate but not the others reduced cellular cholesterol, we studied the ability of butyrate on the rate-limiting enzyme of the cholesterol biogenesis pathway and opted to study the status of HMGCR by western blot in butyrate treated and untreated Huh7 cells. It was observed that there was a significant decrease in HMGCR in butyrate treated cells as compared to untreated cells as a function of butyrate concentration (Fig S1B). The results were presented by densitometric analysis (Fig S1C). The percentage decrease in HMGCR in Huh7 cells upon the addition of 10 mM and 20 mM butyrate were 37 % and 74 %, respectively. Based on the cell line results we asked the same question in the animal model.

### Butyrate corrects hypercholesterolemia in mice receiving HFD (HFD-mice) and reverses elevated hepatic enzymes

The mice were fed with HFD for 30 days and then divided into two groups. One group of mice continued to receive HFD (HFD-mice) and another group received butyrate with HFD for another 15 days (HFD-butyrate-mice). A third group of mice kept as control received regular chow diet (chow-mice). All three groups were sacrificed on day-45 to study a variety of parameters. To start with, body weight and food intake of all three groups of mice were recorded. It was observed that in the HFD-mice the body weight continued to increase at a higher rate as compared to chow-mice up to the 30-day treatment protocol. With the commencement of butyrate treatment on day 31 in the HFD-mice for another 15 days (HFD-butyrate-mice), there was a significant decrease in body weight as compared to HFD-mice. Or in other words, on day 45 there was hardly any difference noticed in body weight between normal and HFD-butyrate-mice (Fig S2A, inset). It was observed that all three groups showed similar magnitude of food intake (Fig S2B). Then we went on to study the status of cholesterol and lipoproteins in all the three groups.

It was observed that compared to chow-mice, HFD-mice showed 60% increase in serum cholesterol which returned to normal in HFD-butyrate-mice (Fig 1). Two liver specific enzymes alanine transaminase (ALT) and aspartate aminotransferase (AST) were studied as the markers of HFD-induced hepatic stress. It was observed that both ALT and AST were significantly increased in HFD-mice which returned to normal in HFD-butyrate-mice (Fig S2C).

**Figure 1:**
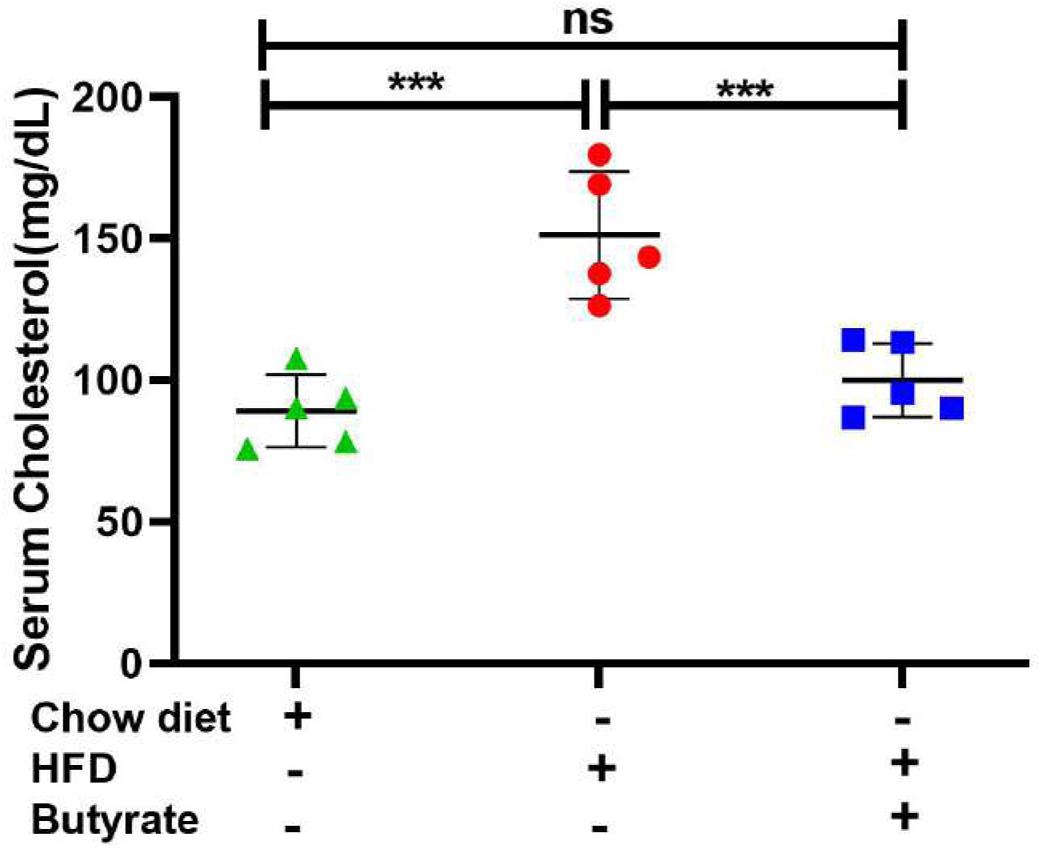
Effect of butyrate treatment on serum cholesterol level of HFD (high fat diet) fed mice (HFD-mice) Female mice (10 in number) were fed with HFD for 30 days and then divided randomly into two groups equally. One group continued receiving HFD (HFD-mice) and another group received 5% sodium butyrate (butyrate) supplemented with HFD (w/w) for another 15 days (HFD-butyrate-mice). Another independent group of mice (5 in number) were fed with normal chow diet for 45 days which served as a control (chow-mice). On day 45 serum cholesterol levels in chow-mice, HFD-mice and butyrate-HFD-mice was determined and expressed in mg/dL. N = 5/ group, experiment was repeated thrice. The data is represented as mean ±SE. ns represents not significant, *** represents p<0.001.

### Butyrate influences cholesterol metabolic genes

The microarray datasets comparing butyrate-treated colonic epithelial cells (MCE301) vs untreated control (GSE4410) and HeLa cells treated with butyrate vs corresponding untreated control (GSE45220) are available in public domain. The results were represented in volcano plots and gene expression showing statistically significant fold change revealed an array of information on cholesterol metabolic genes (Fig S3A). Differential expression analysis indicated that butyrate treatment resulted in an overall repression of cholesterol metabolising genes, of which some are involved directly and some indirectly. The common genes between two data sets which were considered for further studies included HMG CoA reductase *(hmgcr),* acetyl CoA acetyltransferase-2 (*acat2*), 7-dehydrocholesterol reductase *(dhcr7),* HMG CoA synthase 1 *(hmgcs1)* and 24-dehydrocholesterol reductase *(dhcr24*). A significant downregulation in *hmgcr*, *acat2*, *dhcr7*, *hmgcs1* and *dhcr24* expression with butyrate treatment was observed (Fig S3B). By enriching our initial data derived from two independent cell lines, our study was expanded to chow-mice, HFD-mice and HFD-butyrate-mice to study the expression of hepatic cholesterol metabolic genes by qPCR.

### Butyrate treatment to HFD-mice alters cholesterol metabolic gene expression and decreases lipid droplet formation in the liver

Based on the results from cell lines as described in the volcano plot, we studied hepatic expression of *hmgcr, hmgcs*, *acat2* and *dhcr7* by qPCR. There was about a two-fold increase in hepatic *hmgcr* expression in HFD-mice compared to chow-mice, which was returned to normal level in the HFD-butyrate-mice (Fig 2A). It was observed that in the HFD-mice, there was a significant increase in hepatic expression of *hmgcs*, *acat2* and *dhcr7* as compared to the chow-mice, while the expression of only *hmgcs* and *acat2* returned to normal in the HFD-butyrate-mice. The expression of *dhcr7* in the HFD-butyrate-mice decreased significantly compared to the HFD-mice but its expression remained significantly higher compared to the chow-mice. In addition, we opted to study the expression of the cholesterol catabolic enzyme, which did not surface in the Volcano plot: 7A1 of the cytochrome P450 family (*cyp7a1)*. *cyp7a1* expression was significantly reduced in HFD-mice compared to chow-mice and returned to the normal level in HFD-butyrate-mice (Fig 2A).

**Figure 2:**
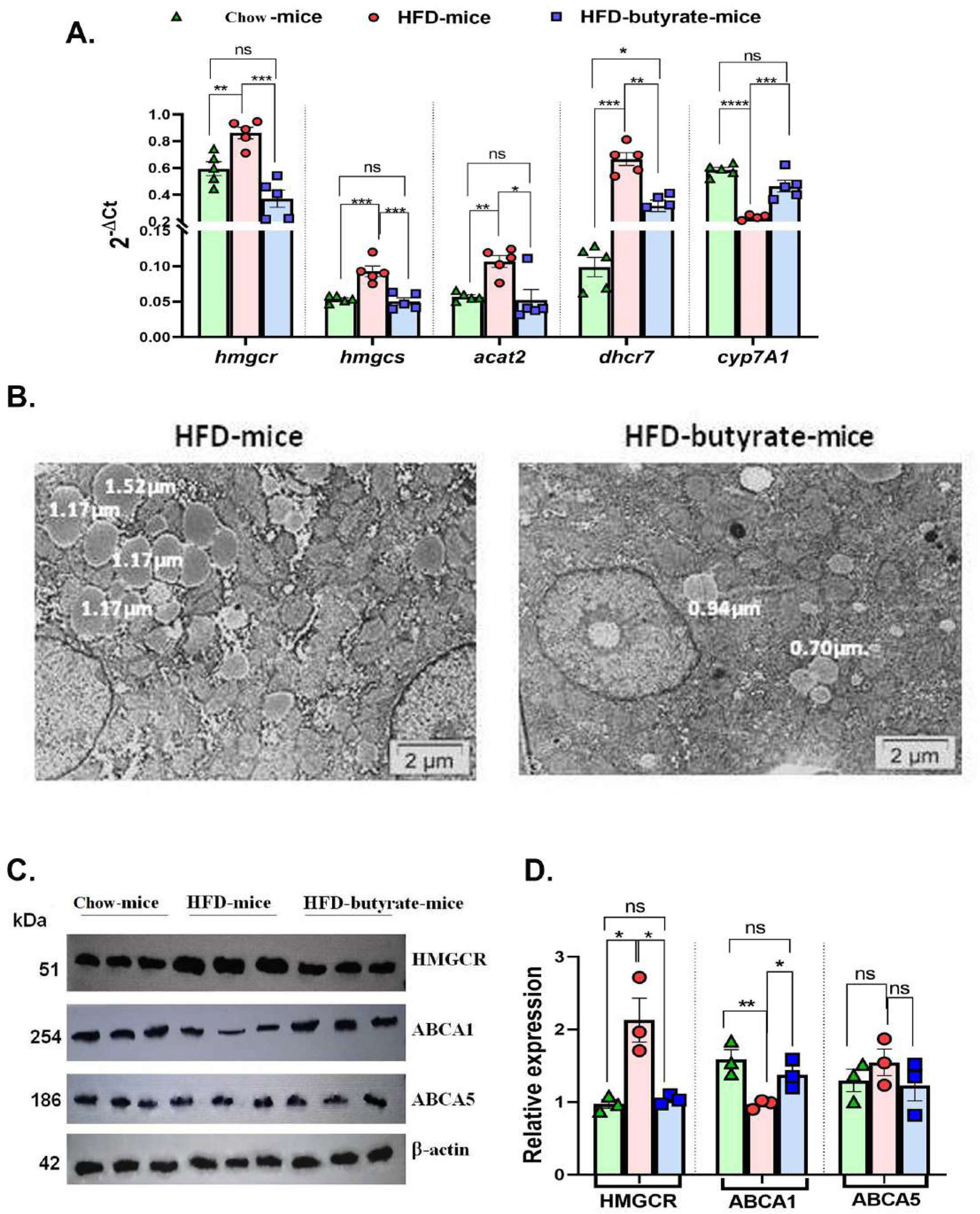
Analysis of hepatic expression of cholesterol metabolic genes by qPCR, lipid droplet status in the liver and HMGCR, ABCA1 and ABCA5 expression by western blot in chow-mice, HFD-mice and HFD-butyrate-mice. Hepatic expression of cholesterol metabolic pathway: *hmgcr*, *hmgcs*, *dhcr7*, *acat-2* and *cyp7A1* was studied in all the groups by qPCR (A). TEM images of the liver sections showing lipid droplets. The sections were visualized under a FEI Tecnai 12 Biotwin transmission electron microscope (FEI, Hillsboro, OR, USA) at an accelerating voltage of 100 kV. Scale bar in the image is 2µm (B). Analysis of hepatic expression of HMGCR, ABCA1 and ABCA5 by western blot (C). Densitometry was done using ImageJ showing relative expression with respect to β-actin control (D). N = 5 / group, data is represented as mean ±SE. The experiment was repeated thrice. *** represents p<0.001, ** represents p<0.01, * represents p<0.05, ns represents not significant.

The liver sections from the mice were inspected by TEM and the representative images of sections were analysed highlighting the preponderance of lipid droplets. It was observed that there was an abundance of lipid droplets of an average diameter of ~1.2 µm in the HFD-mice whereas in HFD-butyrate-mice the droplets were significantly reduced not only in number but also by size with an average diameter of ~0.7 µm (Fig 2B).

Since HMGCR is a key regulatory enzyme of cholesterol biogenesis the results observed in qPCR study were further validated by western blot. It was observed that there was two-fold increase in the expression of HMGCR in HFD-mice compared to chow-mice, while the expression returned to normal in HFD-butyrate-mice (Fig 2C, D).

The hepatic expression of ABCA1 and ABCA5 protein was studied by western blot (Fig 2C). Densitometry analysis revealed that there was a significant decrease in ABCA1 expression in HFD-mice compared to chow-mice, and returned to normal status in the HFD-butyrate-mice (Fig 2D). Notably, the expression of ABCA5 remained unchanged in all the three groups (Fig 2C and D). For further functional analysis of efflux proteins, we used Huh7 cell line.

### Butyrate up-regulates ABCA1 but not ABCA5 expression and increases cholesterol efflux in Huh7 cells

Butyrate treatment of Huh7 cells showed progressive increase in ABCA1 expression as a function of butyrate concentration as evident from the western blot (Fig S4A), and its corresponding densitometry. On the other hand, expression of ABCA5 remained unaltered regardless of butyrate treatment (Fig S4B).

To study efflux, we used 22-NBD-cholesterol-loaded Huh7 cells treated with or without butyrate. The 22-NBD-cholesterol loaded cells were kept for 18 h in medium before butyrate treatment to allow aqueous phase diffusion of cholesterol. It was observed that in the absence of HDL, regardless of butyrate treatment there was negligible efflux of cholesterol. Interestingly, there was a significant increase in cholesterol efflux from 22-NBD-cholesterol loaded Huh7 cells as a function of HDL concentration which was significantly enhanced while cells were pre-treated with butyrate (Fig S4C).

### Butyrate down regulates miR122 biogenesis and upregulates Occludin, a classical target of miR122 in Huh7 cells

We then monitored the pre-miR122, miR122 and Dicer1 status in Huh7 cells by qPCR. It was observed that there was a decrease in miR122 (Fig 3A) and Dicer 1 (Fig 3B) with a concurrent increase in pre-miR122 (Fig 3A) expression as a function of butyrate concentration. We also observed that propionate and acetate at 20 mM did not show any difference in premiR122, miR122 and Dicer1 with respect to control (Fig S4D).

**Figure 3:**
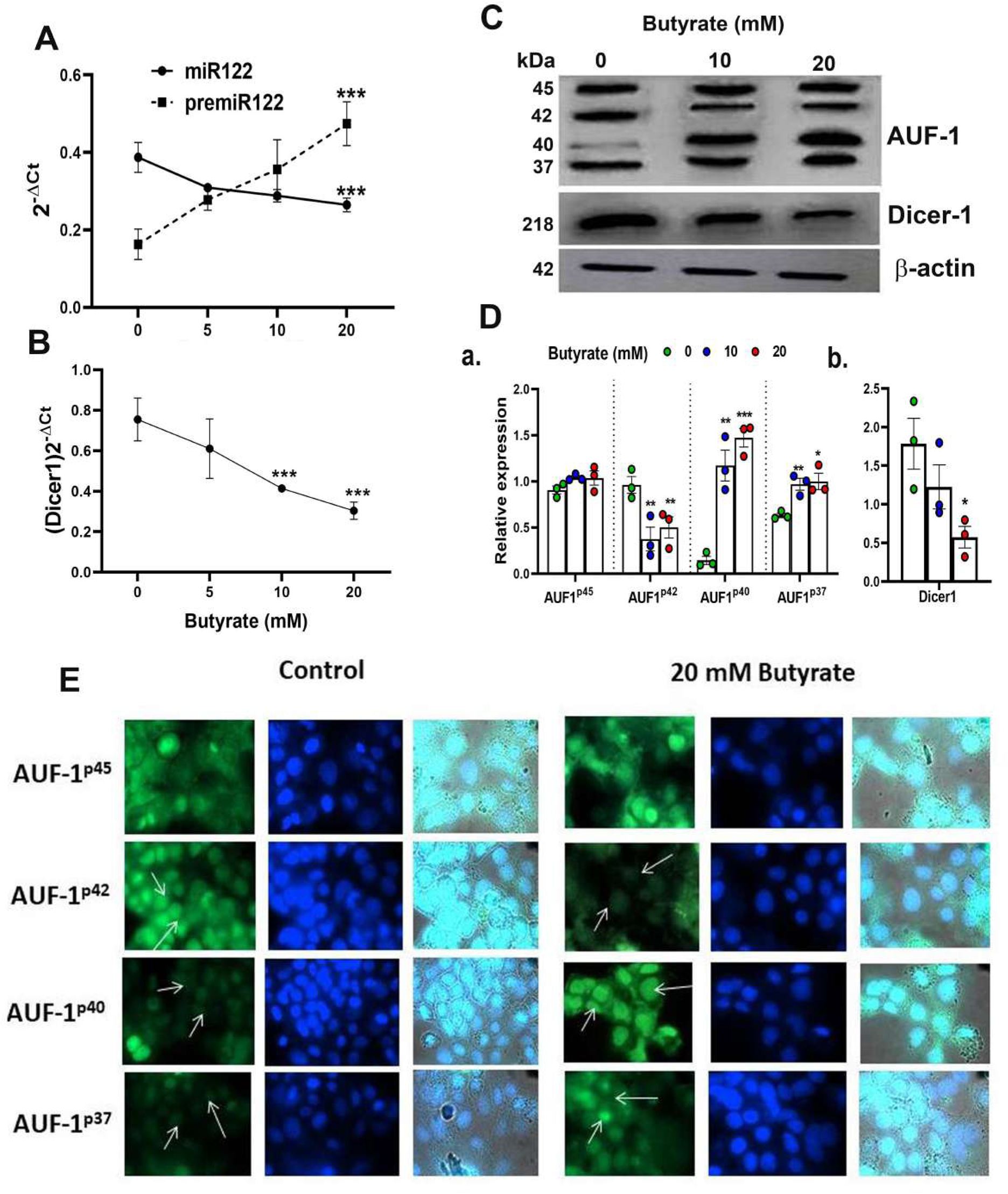
Effect of butyrate on miR122 maturation, Dicer1 expression, AUF1 isoform expression by western blot and EGFP reporter assays in Huh7 cells. Expression of pre-miR122 and miR122 (A) and Dicer1 (B) as determined by qPCR and western blot of AUF1 isoforms (upper panel) and Dicer1 protein (middle panel) as a function of butyrate concentration (C) and corresponding densitometry of AUF1 isoforms (AUF1^p45^/AUF1^p42^/AUF1^p40^/AUF1^p37^) (D, a) and Dicer1 (D, b) with respect to β-actin control using ImageJ. Transfection of Huh7 cells with individual AUF1 isoforms of EGFP reporter construct (pEGFP-AUF1^p45^ or pEGFP-AUF1^p42^ or pEGFP-AUF1^p40^ or pEGFP-AUF1^p37^) in absence and presence of 20mM butyrate for 24 h (E). The resulting fluorescence images are taken under 60X in Apotome Zeiss microscope with a CCD camera controlled with ZEN software (Carl Zeiss, Gottingen, Germany). N=3. The data is represented as mean ± SE. *** represents p<0.001, * represents p<0.05.

We then determined the implication of miR122 deregulation on its target gene expression for which the expression of the tight junction protein (TJ), Occludin was studied which is important to maintain hepatic integrity. It was shown previously that miR122 binds to the 3’UTR of the mRNA of Occludin, causing its degradation ^39^. We show an increase in Occludin expression as a function of butyrate concentration (Fig S5A) demonstrating that butyrate indeed causes a decrease in miR122 as it increases the expression of its target.

### Butyrate upregulates AUF1 and downregulates Dicer1 and Sphingosine kinase 1 (a classical target of AUF1) in Huh7 cells

We studied the expression of the post transcriptional regulator of Dicer1, AUF1, in butyrate-treated Huh7 cells using western blot (Fig 3C). The basal expression of AUF1 isoforms in Huh7 cells was initially p45>p42>p37>p40 which altered to p40>p45>p37>p42 due to butyrate treatment. From the densitometry analyses it was clear that the expression of AUF1^p45^remained unaltered whereas that of AUF1^p42^ significantly decreased due to butyrate treatment with respect to the untreated control. On the other hand, there was an upregulation of the other two isoforms due to butyrate treatment. The fold increase for AUF1^p40^ after the addition of 10 mM and 20 mM of butyrate was 7.36 and 8.7, respectively, with respect to the control. On the other hand, there was a marginal increase in AUF1^p37^ expression after 10 mM butyrate addition, and a low but significant increase after 20 mM of butyrate supplementation (Fig 3D). Furthermore, there was a progressive decrease in Dicer1 as a function of butyrate concentration which was 4-fold in the case of 20 mM butyrate treated Huh7 cells (Fig 3D). We subsequently validated the results by transfection of Huh7 cells with EGFP constructs of each of the AUF1 isoforms in the presence and absence of 20mM butyrate (Fig 3E). We show that butyrate treatment of Huh7 transfected cells resulted in a decrease in EGFP-AUF1^p42^ fluorescence and an increase in EGFP-AUF1^p40^ and EGFP-AUF1^p37^ fluorescence within transfected cells. Conversely, EGFP-AUF1^p45^ fluorescence remained essentially unaltered with or without butyrate treatment. It is also worth mentioning that unlike butyrate, propionate and acetate did not show any significant increase in AUF1^p40^ expression as studied by qPCR (Fig S4D).

Since AUF1 is mRNA decay-promoting protein, for which there may be other important targets other than Dicer1, we also studied the expression of sphingosine kinase (Sphk1). The expression of sphingosine kinase 1 (Sphk1) mRNA, which is another target of AUF1 ^23^ was also decreased upon butyrate treatment. We also observed significant decrease in Sphk1 expression by butyrate treatment to Huh7 cells (Fig S5B). The reciprocal relation of AUF1 and Sphk1 expression upon butyrate treatment offered credence to the notion that butyrate acts by targeting AUF1.

### AUF1 silencing increased Dicer1 and miR122 expression in Huh7 cells

The importance of AUF1 on the expression of Dicer1 and miR122 was studied by silencing AUF1 in Huh7 cells to which the following treatments were added: scramble siRNA (scramble), scramble plus butyrate, siAUF1 alone, or siAUF1 plus butyrate. The expression profile of AUF1 was analysed by western blot (Fig 4A). Interestingly, silencing AUF1 with siAUF1 led to reduction in all the isoforms of AUF1 regardless of the presence or absence of butyrate (Fig 4A). Other cellular consequences due to AUF1 silencing like the expression of Dicer1, miR122 and cellular cholesterol status were also studied. It was observed that silencing with si-AUF1 resulted in significant increase in Dicer1 (Fig 4B), miR122 (Fig 4C) and cellular cholesterol (Fig 4D) regardless of butyrate treatment but this was not seen when the cells were transfected with scramble siRNA. Since the upregulation of AUF1^p40^ was quite prominent of all the AUF1 isoforms due to butyrate treatment, rest of the studies were carried out with the AUF1^p40^ isoform.

**Figure 4:**
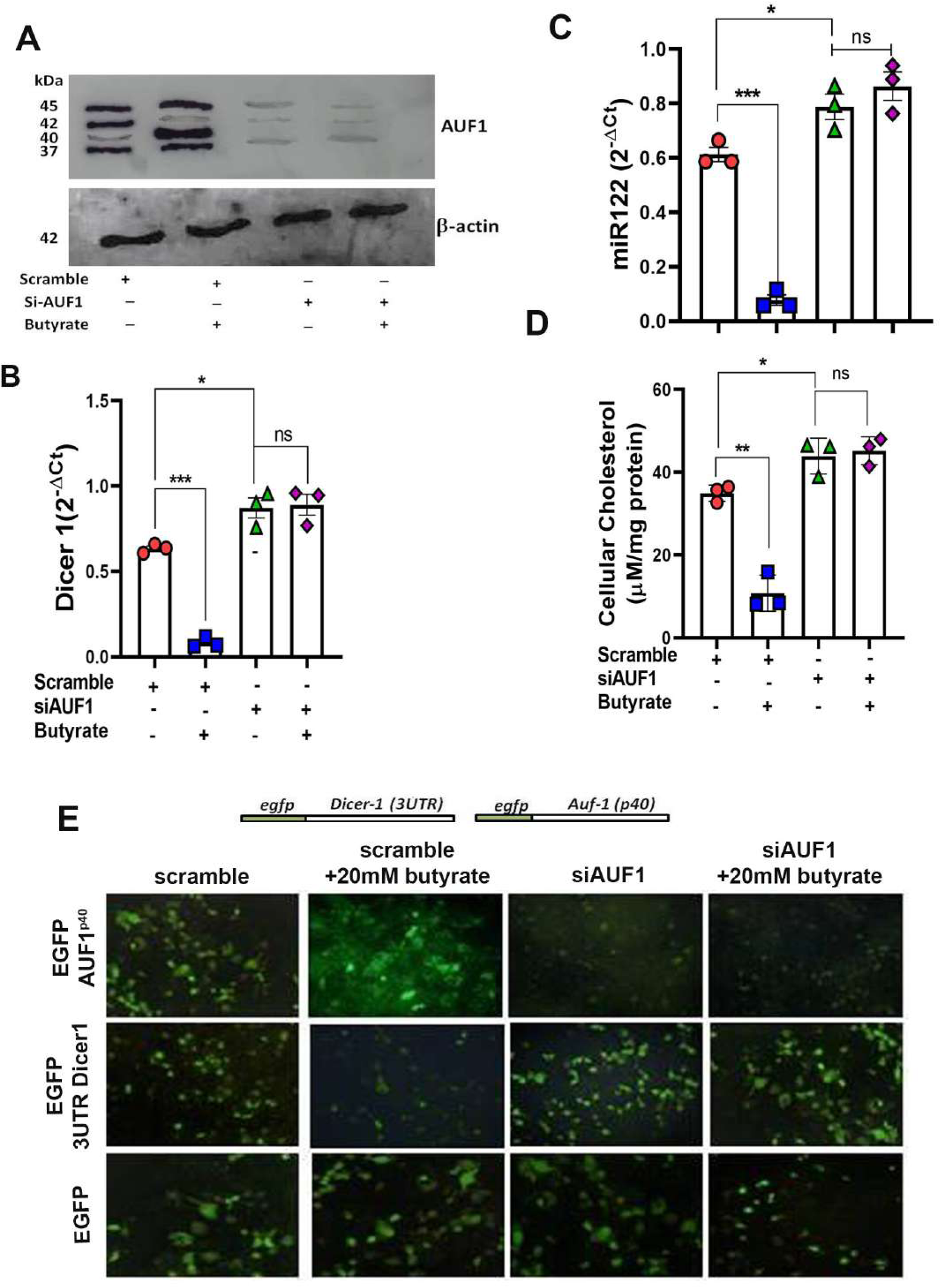
Silencing of AUF1 by siRNA enhanced miR122, Dicer1 expression and cellular cholesterol in Huh7 cells regardless of butyrate treatment. Western blot of AUF1 isoforms after co-transfecting either with siAUF1 or scramble-RNA (scramble) where β-actin used as control (A). The corresponding expression of Dicer1 (B) and miR122 (C) as analyzed by qPCR and cellular cholesterol in cell lysate (D). Analysis of EGFP expression upon transfection with reporter construct of EGFP-AUF1^p40^, EGFP-3ÚTR-Dicer1 and EGFP (from top to bottom) in a co-transfection experiment under following conditions: (i) scramble, (ii) scramble plus 20 mM butyrate, (iii) siAUF1 alone and (iv) siAUF1 plus 20mM butyrate (from left to right) (E). Resulting fluorescence images of co-transfected cells were captured in 20X magnification in Carl Zeiss microscope equipped with a CCD camera controlled with ZEN software (Carl Zeiss, Gottingen, Germany). N=3, the data is represented as mean ± SE. ns represents non-significant, *** represents p<0.001, ** represents p<0.01, * represents p<0.05, ns represents not significant.

To show that AUF1 repressed Dicer1 by binding to the 3’-UTR of its mRNA, a co-transfection experiment was performed. Huh7 cells were transfected either with pEGFP-AUF1^40^ or p-EGFP-3ÚTRDicer1 or pEGFP and the resulting EGFP fluorescence was analysed by fluorescence microscopy. Following the second round of transfection either with scramble or siAUF1, the cells were treated with or without butyrate generating four experimental sets from left to right as follows: (i) scramble, (ii) scramble plus butyrate, (iii) siAUF1 and (iv) siAUF1 plus butyrate. From top to bottom transfection was done with pEGFP-AUF1^40^ or p-EGFP-3ÚTRDicer1 or pEGFP respectively. In the first panel pEGFP-AUF1^p40^, fluorescence was observed when co-transfected with scramble which increased after butyrate treatment. On the other hand, the fluorescence of EGFP-AUF1^p40^ was diminished with siAUF1 and the effect remained the same even after butyrate treatment. In the second panel for the p-EGFP-3ÚTR Dicer1 probe, there was a basal level of fluorescence due to scramble siRNA which diminished after butyrate treatment. The fluorescence of EGFP-3ÚTRDicer1 was increased with siAUF1, but remained essentially unchanged after butyrate treatment. The third panel showed that the EGFP expression remained same during the course of above treatments (Fig 4E).

So far, the relation among butyrate-AUF1-Dicer1-miR122 was studied in Huh7 cells. We further translated these findings in mice model.

### Hepatic expression of miR122, Dicer1 and AUF1^p40^ in chow-mice, HFD-mice and HFD-butyrate-mice

It was observed that there was a significant decrease in pre-miR122 with increase in miR122 in HFD-mice as compared to chow-mice. This expression returned to normal in the HFD-butyrate-mice (Fig 5A, 5B). It was also observed that there was an increase in Dicer1 (Fig 5C) with a concomitant decrease in AUF1^p40^ (Fig 5D) in the HFD-mice and was restored to normal levels in the HFD-butyrate-mice.

**Figure 5:**
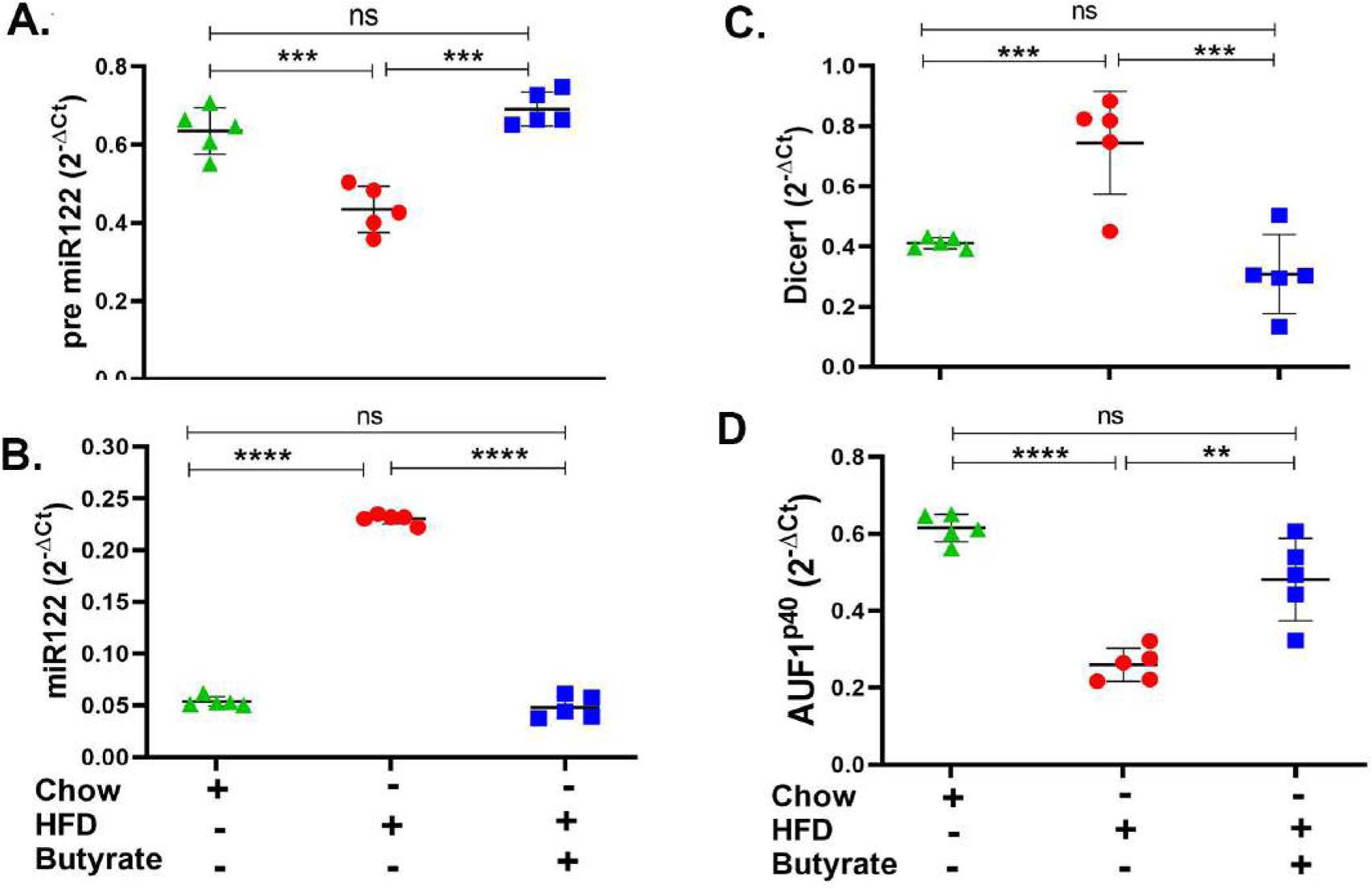
Hepatic miR122, Dicer1 and AUF1^p40^ in chow-mice, HFD-mice and HFD-butyrate-mice. Hepatic expression of pre-miR122 (A), miR122(B), Dicer1(C), and AUF1^p40^ (D), in chow-mice, HFD-mice and HFD-butyrate-mice by qPCR. N=5 / group, data is represented as mean ±SE. The experiment was repeated thrice. ns represents non-significant, **** represents p<0.0001, ***represents p<0.001, ns represents not significant.

### Butyrate downregulates other microRNAs

Since there are reports that mir27 plays an important role in lipid metabolism, we also studied hepatic expression of miR27a and miR27b in chow-mice, HFD-mice and HFD-butyrate-mice. We showed that there was a 6-fold increase in hepatic miR27a expression in HFD-mice compared to chow-mice, which returned to normal levels in the HFD-butyrate mice. Interestingly, miR27b remain unaltered in all the three groups (Fig S6).

### Oral butyrate or probiotic treatment restores gut butyrate-producing bacteria, increases faecal butyrate and reduces serum cholesterol in antibiotic treated mice following AUF1-Dicer1-miR122 pathway

To show that the gut-derived butyrate is intimately linked with cholesterol metabolism in mice, we depleted the gut microbiota of mice by daily gavage supplemented with a cocktail of antibiotics as a surrogate of germ-free mice. The treatment protocol of each group of mice is schematically presented in Fig 6A. The mice were divided into four groups: Group-I: fed with chow diet only (untreated-mice), Group II: received cocktail antibiotic (Abx-mice), Group-III: treated with the cocktail antibiotic for 7 days followed by probiotic treatment every alternate day for 21 days (Abx-probiotic-mice) and Group-IV: Abx-mice receiving butyrate from day 8 to 21 (Abx-butyrate-mice). We determined the relative abundance of the terminal enzyme [butyryl-CoA: acetate CoA-transferase (*butCoAT*)] of the butyrate synthesis pathway from faecal DNA samples as an indirect indicator of the relative abundance of butyrate-producing bacteria. Abx-mice showed a ~100-fold decrease in the relative abundance of *butCoAT* gene compared to untreated-mice which was restored to essentially a normal level in Abx-probiotic-mice (Fig 6B). To show that a decrease in relative abundance of *butCoAT* gene was faithfully reflected in butyrate production, we estimated the faecal butyrate by LC-MS. The decrease in faecal butyrate was more pronounced than the *butCoAT* gene abundance in Abx-mice. There was a ~9000-fold reduction in faecal butyrate in Abx-mice compared to untreated-mice whereas with probiotic treatment, the faecal butyrate increased 70,000-fold higher compared to Abx-mice and ~7 fold high than untreated-mice (Fig 6C). We measured serum and liver cholesterol and showed that the serum cholesterol was increased 2.5-fold on day 7 and remained same on day 21 in the Abx-mice compared to untreated-mice (Fig S8). In Abx-probiotic-mice and Abx-butyrate-mice, the hepatic and serum cholesterol decreased significantly compared to Abx-mice and returned to normal levels (Fig 6D and E).

**Figure 6:**
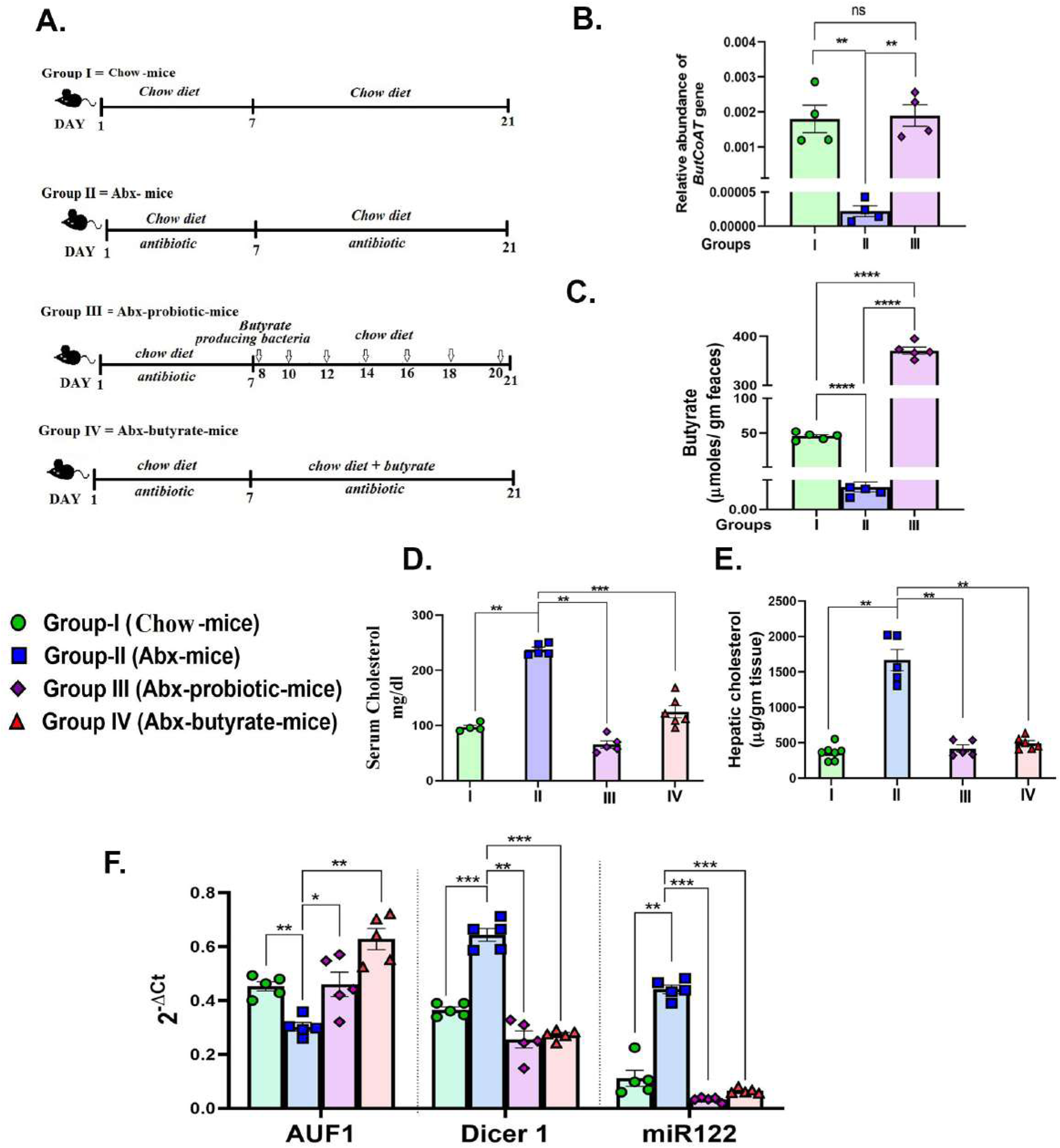
Effect of antibiotic treatment and subsequent probiotic treatment on faecal butyrate, serum and cholesterol, AUF1^p40^, Dicer1 and miR122 expressio. Female mice were used for the study. Experimental design of the study: Group I = untreated chow fed mice (untreated-mice). Group-II = mice receiving cocktail antibiotic and chow diet for 21 days (Abx-mice). Group III= mice receiving cocktail antibiotic for first 7 days followed by bowel cleansing by PEG and administration of probiotics (10^7^ cfu in 200 μl) *via* oral gavage/ day, on every alternate day till 2 weeks (Abx-probiotic-mice). Group IV = Abx-mice fed with 5% butyrate fortified chow diet (w/w) from day 7 to day 21 (Abx-butyrate-mice) (A). Blood, liver tissue and faecal samples were collected on day 21. The relative abundance of *butCoAT* gene in the faeces measured by qPCR and normalized to 16S rRNA bacterial gene abundance (B). Faecal butyrate (expressed in µg/ gm of faeces) on day 21 estimated by LC-MS (C).The serum cholesterol in mg/dL (D) and hepatic cholesterol expressed in µg/ gm liver tissue (E). Expression of hepatic AUF1^p40^, Dicer1 and miR122 determined by qPCR (F). N=5 /group, data is represented as mean ±SE. The experiment was repeated twice. *** represents p<0.001, **represents p<0.01, * represents p<0.05, ns represents non significant.

The rise in serum cholesterol and decrease in faecal butyrate production was correlated with a ~1.7-fold decrease in AUF1 and a concomitant 2-fold and 4-fold increase in Dicer1 and miR122, respectively, in Abx-mice, compared to untreated-mice. As expected, there was significant increase in AUF1 and decrease in Dicer1 and miR122 in Abx-probiotic-mice and Abx-butyrate-mice, compared to Abx-mice (Fig 6F).

### MiR122 overexpression rescues hypo-cholesterolemic effect of butyrate in mice

From the previous experiments it was apparent that butyrate regulates cholesterol biogenesis by exploiting miR122. Therefore, we validated our observation by overexpressing miR122 in butyrate treated mice for which plasmid expressing miR122 was injected through the tail vein of mice fed with chow diet supplemented with butyrate. Another group of mice were injected with mock plasmid and on day-4 post injection, the animals were sacrificed, following which serum and liver cholesterol were analysed. We observed 25-fold increase in miR122 expression in the liver coupled with 4-fold increase in serum cholesterol compared to butyrate-fed mock injected mice (Fig 7 A and B).

**Figure 7:**
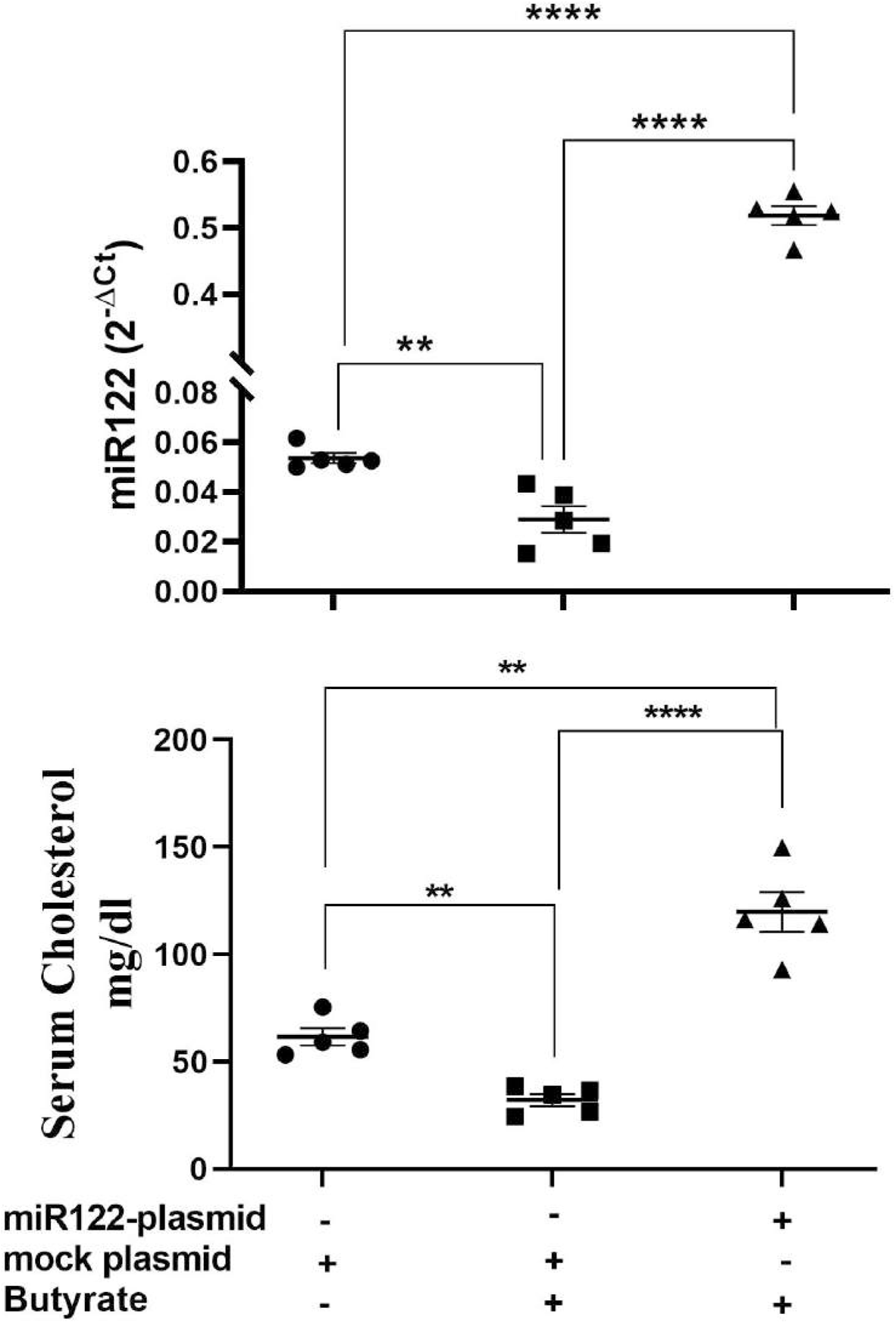
miR122 over expression in liver rescues butyrate induced decrease in serum cholesterol. Each butyrate treated female mice were injected with either 25 µg in 100 µl of miR122 expressing plasmid or 25 µg in 100 µl of mock plasmid in tail vein. The mice were sacrificed 4 days post injection. The serum cholesterol and hepatic miR122 expression was measured. N=5, the data is represented as mean ± SE. The experiment was repeated twice. **** represents p<0.0001, *** represents p<0.001, ** represents p<0.01

### Effect of AUF1 knockdown by Morpholino oligomer (GMO-PMO) on the expression of Dicer1, miR122, cholesterol metabolic enzymes and on serum cholesterol levels

To establish the basic tenet that AUF-1 is an important factor in the butyrate-mediated cholesterol homoeostasis, we opted for AUF1 knock down (AUF1-KD) in mice using novel cell penetrating morpholino oligomer (GMO-PMO). In this study, GMO-PMO specific to AUF1 was denoted as AUF1-MO and corresponding scramble one was denoted as scramble-MO. Mice that received morpholino on day 0 and 7, were sacrificed on day 14, as depicted pictorially (Fig 8A). We showed that there was no difference in AUF-1 expression between normal and scramble-MO treated mice showing that the scramble-MO did not interfere in AUF-1 expression, thereby reinforcing its uniqueness (Fig S10 A, B) and in subsequent experiments normal mice were not included. The results showed that a significant downregulation of the AUF-1 isoform not only in the liver (Fig S10A), but also in other organs such as the kidneys and heart (Fig S10C). Furthermore, butyrate treatment in the scramble-MO or AUF-1 MO mice did not change their respective AUF-1 status in liver (Fig 8B). AUF1-KD in hepatic miR122 and Dicer-1 expression was studied and it was observed that compared to scramble-MO alone, combination of butyrate with scramble-MO led to significant down regulation of miR122 and Dicer-1. As expected AUF1-MO treatment in mice led to upregulation of both miR122 and Dicer-1 and remained unaltered when combined with butyrate treatment (Fig 8C). We also studied the impact of upregulation of miR122 in AUF1-KD mice for which two genes of cholesterol metabolic enzymes, HMGCR and CYP7A1 are considered of cholesterol homeostatic pathway and are the known targets of miR122. As expected, in the scramble-MO group receiving butyrate, there was downregulation of *hmgcr* coupled with up-regulation of *cyp7A1*. In AUF1-KO mice, regardless of butyrate treatment, there was a significant upregulation of *hmgcr* coupled with down regulation of *cyp7A1* (Fig 8D). Both miR122 and Dicer-1 capture a reciprocal relation between *hmgcr* and *cyp7A1* (Fig 8E). As expected, butyrate treatment of scramble-MO-mice showed significant decrease in serum and hepatic cholesterol whereas AUF-1-MO-KD mice showed amply surged cholesterol level which remained unaltered regardless of butyrate treatment (Fig 8E).

**Figure 8:**
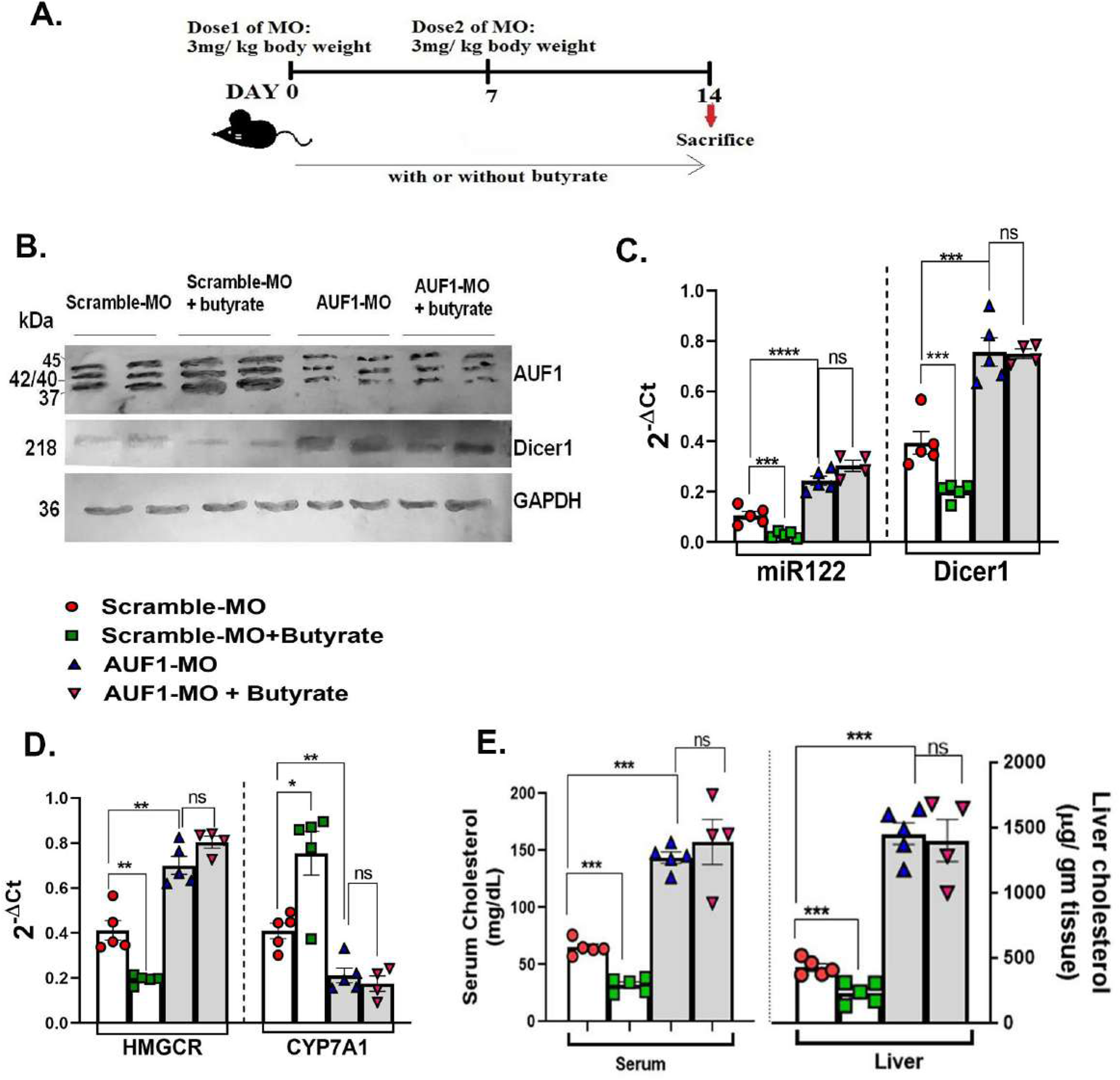
In vivo knock down of AUF1 increases cholesterol synthesis by exploiting AUF1-Dicer1-miR122-cholesterol pathway regardless of butyrate treatment. Female mice (20 in number) were divided randomly into four groups: AUF1-MO injected mice (AUF1-MO-mice), AUF1-MO injected mice fed with butyrate supplement (butyrate-AUF1-MO-mice), scramble-MO injected mice (scramble-MO-mice) and scramble-MO injected mice fed with butyrate supplement (butyrate-scramble MO-mice) the scheme as shown schematically (A). Western blots to determine hepatic expression of AUF1 and Dicer1 (B). GAPDH was used as control. AP-conjugated secondary antibodies were used in western blot. Expression of hepatic miR122 and Dicer1 determined by qPCR (C). Hepatic expression of cholesterol metabolising enzymes: HMGCR and Cyp7A1 determined by qPCR (D). Serum and liver cholesterol was measured and expressed in mg/dL and µg/gm tissue respectively (E). N=5/ group, data is represented as mean ±SE. The experiment was repeated twice. ns represents non-significant, **** represents p<0.0001, ***represents p<0.001, ns represents not significant.

## Discussion

This discourse attempts to shed light on a deeper understanding of the role of butyrate in host cholesterol homeostasis for which following questions were asked: (a) what are the target genes and the intracellular players for linking effect of butyrate on cholesterol metabolism. (b) whether endogenous gut derived butyrate exploits the similar molecular pathway like exogenously fed butyrate and, (c) what are the phenotype of cell line and mouse when selective important players were either overexpressed like miR122 or knocked down like AUF-1 to bring strength to our notion.

We showed that unlike acetate and propionate, butyrate has emerged as a hypo-cholesterolemic short chain fatty acid both in cell lines and in HFD-mice, which has warranted a mechanistic look to understand how butyrate influences a diverse repertoire of functions. To address the first point we sought assistance of the published microarray datasets (GSE4410 and GSE45220) and showed that the key cholesterol-metabolic genes, *hmgcr*, *hmgcs1, acat2, dhcr7* and *dhcr24* were down regulated upon butyrate treatment which we have also validated by qPCR in HFD-mice and HFD-butyrate-mice except dhcr24, as it is involved in the later stages of cholesterol biosynthesis. The enzymes HMGCR and HMGCS are rate-limiting ^40^ and whereas DHCR7 is the terminal enzyme in the cholesterol synthesis pathway ^41^, while ACAT2 is involved in cholesterol ester formation ^42^. In addition, we also studied the expression of two other important entities: (a) *cyp7A1,* a member of the monooxygenase cytochrome P450 superfamily that catalyses cholesterol to bile acid ^43^ and (b) ABCA1, involved in cholesterol efflux ^11^. The expression of *acat2* and *cyp7a1* appeared to be quite opposite in HFD-mice as compared to HFD-butyrate mice which may have contributed to the ‘lipid droplet’ formation and butyrate treatment may have caused their dissolution. The HFD-mice showed elevated ALT and AST indicating hepatic stress which returned to normal in HFD-butyrate-mice. Our results are concordant with earlier reports on the beneficial effect of butyrate on liver damage and dyslipidemia ^38, 44^.

It has been reported that butyrate induces profound changes in miRNA in MDCK cells^45^. We identified few important miRNA, in particular, those that are associated with cholesterol metabolism like miR122, miR27a and miR27b ^12 13^. In an elegant study by others showing that the treatment of mice with antagomir-miR122 led to the downregulation of a large number of genes, of which the top-ranking functional category includes the cholesterol biosynthesis genes namely, *hmgcr, hmgcs1* and *dhcr7* ^12^. Additionally, miR122 destabilizes cyp7A1mRNA by binding to its 3’UTR ^46^. Reports show that the silencing of miR-122 in African green monkeys and chimpanzees have resulted in substantial reduction in total plasma cholesterol ^47^ while in the Huh7 cell line, it rescues excess lipid deposition 48.

To witness the effect of butyrate on the cholesterol metabolic landscape, we undertook studies with Huh7 cell line, which is known to have higher miR122 expression^49^. The butyrate concentration used was in tune with an earlier study ^38^. For in vitro experiments on a cancer cell like Huh7 relatively higher dose of butyrate was needed for gene expression study. As epigenetic modifier, butyrate may be differentially utilized in normal and cancerous cell^50^, therefore it is tempting to speculate that higher dose of butyrate is required to induce effect in Huh7. Butyrate treatment showed an increase in ABCA1 expression whose function was validated by showing as increase in cholesterol efflux and decrease in miR27a which caused decay of ABCA1 mRNA^13^. To link butyrate with miR122 biogenesis, butyrate-treated Huh7 cells showed reduced Dicer1 expression which halted maturation of miR122 from its precursor. The effect was butyrate-specific, because neither propionate nor acetate caused any effect. We showed an increase in occludin expression upon butyrate treatment of Huh7 cells indicating that butyrate may impart an effective tight junction function. It is reported that miR122 binds to the 3’UTR of the occludin mRNA, causing its decay ^39^. The impaired occludin expression has frequently been associated with chronic liver diseases ^51^.

Since Dicer1 is an important protein for the canonical miRNA biogenesis, the question arises whether all microRNA synthesis is reduced due to Dicer1 downregulation by butyrate. We show that the expression of miRNAs that decay ABCA1 mRNA by binding to its 3’UTR^13^ like miR27a but not miR27b, was inhibited due to butyrate treatment suggesting that all miRNAs were not equally affected due to butyrate treatment as there are reports of Dicer-1 independent miRNA biogenesis ^52–54^ where slicer activity of Argonaute2 plays vital role in pre-miRNA cleavage ^52^. Dicer is critical for most miRNAs, but the 5p miRNAs appear to be produced to an extent without Dicer ^53^.

Cellular steady state mRNA levels are maintained by the result of two opposite mechanisms, i.e., transcription and decay. There is a report that AUF1, a RNA binding protein lowers the Dicer1 mRNA stability by binding to its 3’UTR^19^. The decay of transcripts containing AU-rich elements (AREs) is another example of RNA regulation that coordinates several physiological processes through post-transcriptional regulation^55^. AUF1 is having multitude of functions, such as DNA-binding^56^, RNA turnover^57^ and mRNA translation efficiency^58^. The AUF1 family contains four isoforms AUF1^p45^, AUF1^p42^, AUF1^p40^, and AUF1^p37^ and their relative levels, rather than the absolute amount of individual AUF1 isoforms, determines the net mRNA stability of ARE-containing transcripts ^59^. Although different AUF1 isoforms have different binding affinity to specific mRNA 3’-UTRs ^60, 61^, the detailed mechanism by which AUF1 isoforms selectively regulate Dicer1-mRNA turnover is unclear.

We showed that butyrate induces upregulation of both AUF1^p40^ and AUF1^p37^ but not AUF1^p42^ and AUF1^p45^, in Huh7 cells. Since AUF1^p40^ expression was quite prominent among all the isoforms, rest of the studies were done with the AUF1^p40^ isoform only. But it does not exclude the possibility of the role of AUF1^p37^ in the regulatory processes. There is a previous report stating that butyrate influences alternative splicing of many proteins like vascular endothelial growth factor (VEGF), IL-18, Defensin β-1 gene and some others^62^. Therefore, one may speculate that butyrate, by influencing alternative splicing of AUF1, may regulate gene expression. For functional validation of butyrate-induced upregulation of AUF1, we studied hepatic expression of a known classic target of AUF1 sphingosine kinase1 (spkh1) ^23^, responsible for cellular proliferation ^23^ and prognostic marker for hepatocellular carcinoma (HCC)^63^. Butyrate-mediated decay of Sphk1 may also explain anticancer effects of butyrate on HCC^64^. The precise mechanism by which butyrate activates AUF1 is not clear. A previous study showed that HDAC inhibitor promotes transcription of RNA binding protein (RBP) through the activation of the transcription factors of early growth response protein ^65^. Butyrate being an HDAC inhibitor ^66^ may also activate AUF1 through these transcription factors.

We showed that silencing of AUF1 by siRNA led to down-regulation of all the isoforms of AUF1, coupled with an increase in Dicer1, miR122 and cellular cholesterol, regardless of absence or presence of butyrate. Thus, butyrate may serve as a master regulator of Dicer1 stability through AUF1. The limitation of our study is that the commercially-available siRNA appears to downregulate all the isoforms of AUF1 which makes it is difficult to ascertain the importance of any specific isoform in the process. The above findings portray the sequential interplay of several intracellular players which operate seamlessly as “butyrate-AUF1-Dicer1-miRNA122-cholesterol-metabolic enzymes-cholesterol level” whose elegance was further verified using the HFD induced gut dysbiosis model. A recent study demonstrated that butyrate decreases lipid profile including cholesterol largely by LKB1-AMK-Insig signaling pathway ^38^. Interestingly, miR122 knockdown in mice showed activation of LKB1-AMK pathway ^48^. This study leads credence to the notion that LKB1-AMK-Insig pathway that are shown to connect butyrate and cholesterol converge in the downstream of “AUF1-Dicer1-miRNA122” axis.

To address the second query and to show that gut derived butyrate is the prime regulator of cholesterol homeostasis, we depleted gut bacteria by antibiotic treatment as a surrogate of germfree mice ^67^ and then reconstituted with probiotics. The probiotic consisted of *Clostridium butyricum*, lactic acid-producing bacteria (LAB), *Bacillus mesentericus* and *Streptococcus faecalis* which produce SCFA. *Clostridium butyricum* is a butyrate producer ^68^, LAB produces lactate which is fermented to butyrate ^69^, *Streptococcus faecalis* produces SCFAs ^70^ while *Bacillus mesentericus* prevents the growth of pathobionts and increases the abundance of Bifidobacteria (SCFA producing bacteria) ^71^.

The final step of butyrate production is usually catalysed by *butCoAT* and butyrate kinase (*buk*) ^72^. In a metagenome-based study, it was predicted that the *butCoAT*-mediated route is 10-fold more abundant than that mediated by *buk* ^73^. In our investigation, we have studied the butyrate production indirectly by measuring the relative abundance of *butCoAT* gene in the faecal samples, and directly by measuring the faecal butyrate concentration by LC-MS. Oddly enough, we observed that the faecal butyrate concentration was much higher as compared to *butCoAT* gene abundance in the probiotic treated group; the cause of such temporal mismatch is not clear. Usually, the acetyl CoA pathway contributes to 79.7 % of all butyrate production in gut microbes ^74^. But other pathways like lysine, glutarate, and 4-aminobutyrate pathways also contribute to butyrate production ^74^. These pathways are prevalent in Firmicutes and some other phyla, such as Fusobacteria and Bacteroidetes ^75, 76^. It is tempting to speculate that other butyrate-producing pathways may operate to contribute to the total butyrate pool. Harmonious to our result of increased serum and liver cholesterol in Abx-mice, earlier report showed that depleting intestinal microbiota by antibiotics enhanced bile acid absorption and expression of HMGCR and HMGCS in liver resulting in 55% increase in serum cholesterol ^77^. Another study showed that germ-free mice display 1.5 times higher liver cholesterol than conventional mice ^78^.

To address the third point, we showed that mir-122 is indeed involved in the recovery of serum cholesterol, we over expressed miR-122 in butyrate-treated mice which showed appreciable recovery of serum cholesterol. To further establish the link between AUF-1 and cholesterol homeostasis, we knocked down AUF1 by deploying morpholino oligomer (MO) which are short single-stranded DNA analogues that are built upon a backbone of morpholine rings and are resistant to host enzymes present, a characteristic that makes them highly suitable for *in vivo* applications. In MO, the guanidinium units that are tethered with the antisense, are more rigid which help to get a favourable conformation to bind with the complementary mRNA ^79^. Lower molecular weight of MO helps in endosomal escape efficiency ^80^. Notably, morpholino-based therapy for Duchenne muscular dystrophy (DMD) has been approved by FDA which is now a hallmark for morpholino-based antisense therapy ^81^. The chimera of GMO and PMO producing MO used in this investigation bears unique testimony of self-transfecting ability and no need of vehicle for delivery, eliminating the possibility of vehicle induced toxicity (doi: https://doi.org/10.1101/2021.06.04.447039). Since AUF1 knock out mice has high mortality due to high septicaemia ^22^, the selective AUF1 knock down (KD) by MO offered unique platform to test our hypothesis *in vivo*. The rise in serum cholesterol along with increase in *hmgcr* and decrease in *cyp7A1* expression in the liver of AUF1 KD mice compared to scramble was noted, indicating that AUF1 is the master regulator of cholesterol biogenesis. Furthermore, it was also observed that exogenous butyrate fails to correct hypercholesterolemia in AUF1 KD mice. Importantly, AUF1 KD showed high levels of Dicer1 and miR122 expression compared to scramble. The increase in miR122 and decrease in *cyp7A1* in AUF1-MO mice is significant as miR122 directly binds to cyp7A1 mRNA ^46^ leading to its decay.

Overall, the present study demonstrates the effectiveness of butyrate as a powerful regulator of cholesterol homeostasis at multiple layers. By aligning series of in vitro and in vivo experiments, we report how the intestinal microbial butyrate regulates cholesterol balance by involving intercellular players in tandem as follows: ‘butyrate-AUF1-Dicer1-miR122-cholesterol metabolic enzymes-cholesterol level where the individual members contribute either via up or down regulation.

## Acknowledgements

We acknowledge the support of Director, ICMR-NICED, Kolkata for carrying out the study. Our deepest gratitude to Prof Syamal Roy (CSIR-IICB, Kolkata) for his support, helpful discussion and suggestions for the manuscript. We thank Ms. Shatarupa Bhattacharya and Rahul Gajbhiye for LC-MS analysis. We acknowledge Dr. Amit Ghosh and Dr. Santasabuj Das (ICMR-NICED) for critically reviewing the manuscript. OD is recipient of fellowship from the DST. MC and AM is recipient of fellowship from CSIR.

## SUPPLEMENTAL INFORMATION

### Materials and Methods

#### Reagents and Chemicals

Chow diet (Harlan Teklad LM-485), High Fat Diet (HFD) (Harlan Teklad TD93075) was purchased from ICMR-NIN, Hyderabad, India. Dulbecco’s modified Eagle’s medium (DMEM) and foetal calf serum (FCS) were purchased from GIBCO (Waltham, MA, USA). Gentamycin, BCA protein assay kit, BCIP-NBT kit were purchased from Thermofisher (Waltham, MA, USA). Penicillin, streptomycin, ampicillin, vancomycin, metronidazole, MTT (3-(4,5-dimethylthiazol-2-yl)-2,5-diphenyl tetrazolium bromide), Triton X100, PMSF, leupeptin, glycine, acrylamide, bis-acrylamide para-formaldehyde, glutaraldehyde, sodium butyrate, sodium propionate, sodium acetate, Hoechst 33342, and cholesterol estimation kit were purchased from Sigma (St. Louis, MO, USA). AST and ALT estimation kit was purchased from Transasia Biomedicals (Mumbai, India). 22-NBD-[22-(*N*-(7-Nitrobenz-2-Oxa-1,3-Diazol-4-yl)Amino)-23,24-Bisnor-5-Cholen-3β-Ol]-cholesterol, cholesterol and phosphatidyl choline were purchased from Avanti-polar lipids (Birmingham, USA). HDL was purchased from Beacon Diagnostics Pvt Ltd (Navsari, Gujrat, India). Probiotic was purchased from Bifilac (Tablets India, Chennai, India). Poly ethylene glycol (PEG) solution was purchased from Jhaver Centre (Chennai, India). Amplex red cholesterol assay kit, Lipofectamine 2000, PVDF membrane, Trizol, Opti-MeM, were purchased from Invitrogen (Carlsbad, CA, USA). QIAamp stool mini kit was purchased from Qiagen (Hilden, Germany). Prime script first strand cDNA synthesis kit, TB Green Premix ex-Taq (Tli RNase H+) qPCR kit were purchased from Takara (Shiga, Japan). (Ripa lysis buffer, anti-β-actin antibody (polyclonal), siAUF1, Anti-AUF1 antibody (rabbit polyclonal), anti-HMGCR antibody (polyclonal), anti-ABCA1 antibody (rabbit monoclonal) were purchased from Cell Signalling Technology (Danvers, MA, USA). Anti-ABCA-5 antibody (rabbit, polyclonal) was purchased from Abcam (Cambridge, UK). Anti-Dicer1 antibody (mouse monoclonal IgG2a) and Anti-AUF1 antibody (mouse monoclonal IgG1) was purchased from Santa-cruz Biotechnology (Dallas, Texas, USA). Anti-GAPDH antibody (rabbit polyclonal) was purchased from Bio-Bharati (Kolkata, India). All primers were purchased from IDT (Lowa, USA). Human hepato-carcinoma cell line Huh7 and pmiR122 were a kind gift from Dr. Suvendranath Bhattacharya (CSIR-IICB, India). pEGFP-3ÚTRDicer1 and pEGFP were a kind gift from Dr. Myriam Goropse, NIH, USA. pEGFP-AUF1isoforms were also gifted by Prof. Andrea Pautz, University of Mainz, Gutenberg.

#### In silico analysis of global microarray data

Raw microarray expression data was obtained in a CEL file format using GSEOquery package (DOI: <u>10.18129/B9.bioc.GEOquery</u>) from GSE45220 (human) and GSE4410 (mice) datasets. The *GEOquery* R package parses GEO data into R data structures that can be used by other R packages. Normalization and background correction of Gene Expression Measurements was performed using oligo package’s Robust Multichip Average (RMA) algorithm that attempts to remove local biases across samples in order to enable relevant differential expression testing. Variations are scanned across the samples using principal component analysis and Hieracrchial clustering method. Following which, differential gene analysis was done using Limma package. The *limma* (Linear Models for Microarray Analysis) R package has evolved as the most widely used statistical tests for identifying differentially expressed genes using Limma based adjusted P-value. P-value of <0.05 are selected. Results are annotated using library hugene10sttranscriptcluster.db for GSE45220, and huex10sttranscriptcluster.db for GSE4410. In case of genes with two or more probes aligned/mapped, only the most significant ones were chosen and the list of upregulated and downregulated genes are prepared accordingly. Genes with the smallest adjusted P-value and highest logFC values are the most significant. Amongst them genes which have a possible role in cholesterol metabolism were selected using KEGG pathway, BioGPS and Reactome databases and were sorted accordingly. All the cholesterol genes showed a downregulated expression. The data were represented as Volcano plot.

#### Transmission Electron Microscopy

Liver tissue was fixed with 3% glutaraldehyde in 0.1M sodium cacodylate buffer. Subsequently, a secondary fixation was conducted with 1% Osmium tetroxide, followed by dehydration with ascending grades of acetone, and finally embedded in Agar 100 resin and polymerization at 60°C. The ultrathin sections (40–50 nm) of the tissue were obtained using a Leica Ultracut UCT ultramicrotome (Leica Microsystems, Germany), picked up on nickel grids, and dual-stained with 2% aqueous uranyl acetate and 0.2% lead citrate. The sections were visualized under a FEI Tecnai 12 Biotwin transmission electron microscope (FEI, Hillsboro, OR, USA) at an accelerating voltage of 100 kV (1).

#### Food consumption

Food was given between 13:00~13:30 p.m. each day to avoid disturbance of the circadian clock. The daily consumption of food in mice was recorded every day by weighing the food given (food placed in the food receptacle at time zero) and food remaining (food left in the food receptacle 24 hours later). The cumulative food intake by each group (5 mice / group) per day was determined.

#### Collection of blood, preparation of serum samples, biochemical analysis, estimation of cholesterol and lipoproteins and hepatic enzymes

Blood collected from tail vein was allowed to stand for 3 h at room temperature and then serum was prepared by centrifugation at 1800 rpm analyzed for serum cholesterol by Assay Kit. Levels of aspartate aminotransferase (AST) and alanine aminotransferase (ALT) in serum were measured in using kit from Transasia Biomedicals (India).

#### Transfection

Cells were plated in 6-well culture plates at the density of 1×106 cells/well and cultured in 2 ml serum-free medium for 24 h to 80% confluence. Transfection was performed with Lipofectamine 2000 according to the protocol recommended by the manufacturer. Briefly, 25 μl Opti-MEM medium was used to dilute 1.0 μl lipofectamine and 0.5 μg plasmid or 0.27 μg siRNA, and equal volume of the plasmid or siRNA and lipofectamine was mixed at room temperature for 15 min. Cells were transfected by adding 100 μl of the Opti-MEM medium containing lipofectamine and plasmid or siRNA at 37° C for 6 h, and then cells were grown at DMEM containing 10% FCS (2).

#### Fluorescence microscopy

Cells grown on glass cover slips were transfected with either pEGFP-AUF1^p40^ plasmid (3) or pEGFP-Dicer-1-3′UTR or p-EGFP (4) for 24 h. After washing with PBS, the cells were treated with 1 µg/ml Hoechst 33342 for 5 min at room temperature and washed again with PBS three times. Fluorescence images were captured with Carl Zeiss microscope equipped with a CCD camera controlled with ZEN software (Carl Zeiss, Gottingen, Germany).

#### Tissue homogenisation, preparation of RNA and Protein

Liver samples were dissected into small pieces and were resuspended either in RIPA Lysis buffer (20mM Tris-HCl pH 7.5, 150 mM NaCl, 1mM EDTA, 1mM EGTA, 1% NP-40, 1% Sodium deoxycholate, 2.5 mM Sodium Pyrophosphate, 1mM β-glycophosphate, 1mM Na_3_VO_4_, 1 µg/ml leupeptin with 1mM of PMSF immediately before use) for protein isolation or in Trizol (Invitrogen, US) for RNA isolation. The tissue was homogenized using a micropestle and centrifuged at 13,000 g for 15 min at 4°C. The clear supernatant was collected and either stored as protein lysate in −80°C or further processed to isolate RNA using the standard protocol (2).

#### RNA extraction and reverse transcription

Cells were cultured in 24-well plates to 80% confluence. Total RNA from cells or tissue was extracted with Trizol according to the protocol recommended by the manufacturer. The concentration of the extracted RNA was analyzed by Nanodrop spectrophotometer (Thermo) and RNA was stored at –80° C. cDNA was prepared from total RNA by reverse specific primers using Super Reverse Transcriptase MuLV Kit. The primers for the reverse transcription are listed in Table 1. U6 and GAPDH were normalized for the expressions of miRNAs and other genes of interest respectively. The total reaction volume for reverse transcription was 20 μl in which 1 μM of reverse primer, 5 ng of RNA template, 1 μl dNTP mix, 12 μl of DEPC treated water, 4 μl of 5X first strand buffer, 1 μl of 0.1 M DTT, 1 μl of RNase inhibitor and 1 μl Super RT MuLV. Reverse transcription was carried out for 65°C for 5 minutes, followed by incubation at 55°C for 1 hour and then heat inactivating the reaction at 70°C for 15 minutes (4).

#### Quantitative real-time PCR

The miRNAs and mRNA levels were quantified with Applied Biosystems^TM^ StepOne^TM^ Real Time PCR System with RT^2^ SYBR^®^ Green qPCR Mastermix following the manufacturer’s instructions. Each 20 μl qPCR reaction contained an amount of cDNA equivalent to 5 ng of total RNA, 10 μl of RT^2^ SYBR^®^ Green qPCR Mastermix, 1 μM of the forward and reverse primer (each) and nuclease free water (5). Real-time PCR was performed with the following conditions: 95°C for 10 min, 40 cycles of 95°C for 30 sec, 60°C for 1 min and 72°C for 1 min PCR product was calculated according to the 2^^^–ΔCt method described previously (4).

The stem loop primer stock was overlaid with 100µl molecular biology grade mineral oil (Sigma). The mixture was heated to 95°C and were kept at 75°C, 68°C, 65°C, 62°C and 60°C for an hour each respectively. Thereafter, working stock of 10µM was prepared and stored in −20°C till further use (6).

#### Primers

**Table S1:**
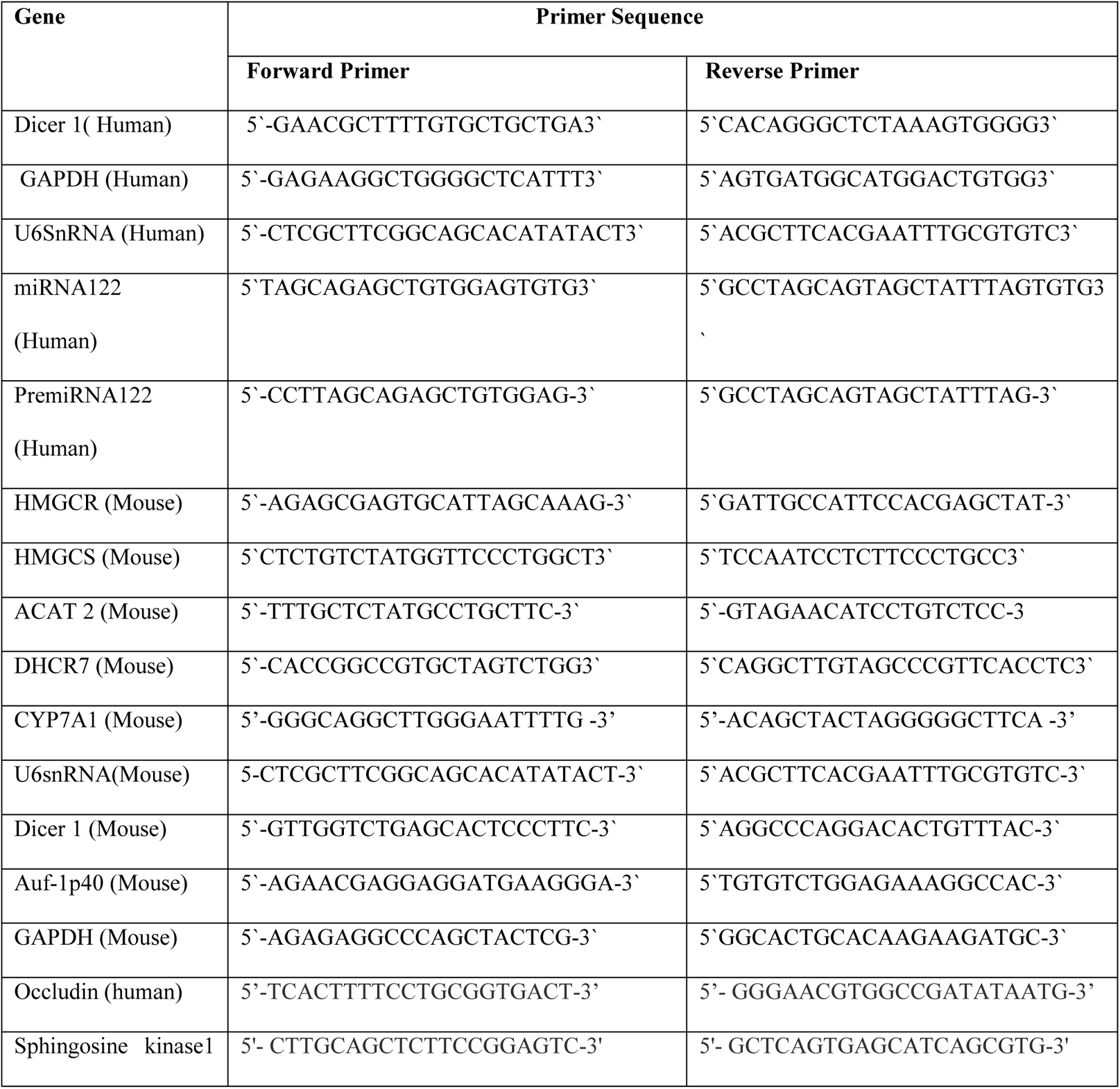
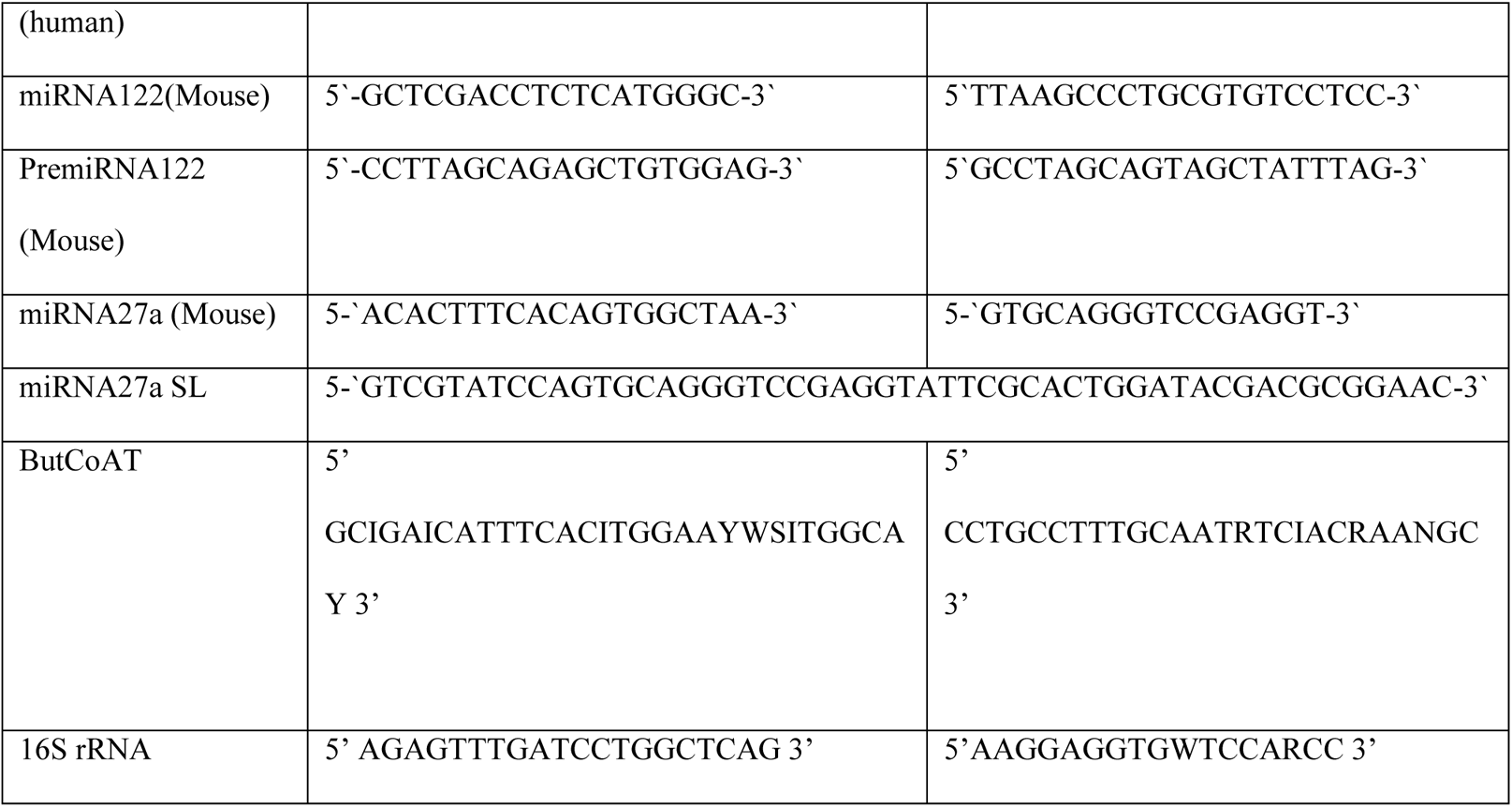
The primer sequences use for PCR amplification are as follows:

#### Western blot

Liver tissue protein or cell lysate were extracted in RIPA Lysis buffer. Protein concentration was measured using Pierce ^TM^ BCA Protein Assay Kit. Proteins (50 μg/lane) were separated by using SDS-PAGE on 10% gel under reducing condition and electro transferred to PVDF membrane in a transferred buffer (25mM Tris-HCl, 150mM Glycine, 20% Methanol). Membranes were blocked at room temperature with 5% non fat skim milk in TBS for 2 hours, and then incubated with primary antibody against specific protein. The membranes were incubated either with the horseradish peroxidase (HRP)-conjugated secondary antibodies or Alkaline phosphatase (AP)-conjugated antibodies at 37° C for 1 h. For HRP-conjugated antibody treatment, SuperSignal West Pico chemiluminescent substrate kit (Thermo) was used to visualize the blotting results. The blots were imaged with Fluor Chem R system (ProteinSimple, San Jose, CA, USA) (4). For AP-conjugated antibody treatment, BCIP-NBT substrate kit (Thermo) was used to visualize the blotting results (7).

**Table S2:** Antibody used for Western Blots

| **Name of Antigen** | **Raised in** | **Source** | **Dilutions used** |
| --- | --- | --- | --- |
| HMGCR | Rabbit | Cell Signalling Technology | 1:1000 |
| ABCA1 | Rabbit | Cell Signalling Technology | 1:1000 |
| ABCA5 | Rabbit | Abcam | 1:1000 |
| AUF-1 (p37 specific) | Rabbit | Cell Signalling Technology | 1:1000 |
| AUF-1/ HNRNP | Mouse | Santa Cruz | 1:100 |
| Dicer 1 | Mouse | Santa Cruz technology | 1:1000 |
| $\beta$ Actin | Rabbit | Cell Signalling Technology | 1:1000 |
| GAPDH | Rabbit | BioBharati | 1:1000 |
| Anti-mouse-HRP<br>conjugated secondary<br>antibody | Horse | Cell Signalling Technology | 1:5000 |
| Anti-rabbit- HRP<br>conjugated secondary<br>antibody | Goat | Cell Signalling Technology | 1:5000 |
| Anti-mouse-AP<br>conjugated secondary<br>antibody | Goat | Abcam | 1:5000 |
| Anti-rabbit-AP<br>conjugated secondary<br>antibody | Goat | Abcam | 1:5000 |

#### Synthesis of Guanidinium Morpholino Oligonucleotides-Protected chlorophosphoramidate Morpholino Oligonucleotides (GMO-PMO)

All reagents were purchased from commercial sources and used without further purification, unless otherwise mentioned. All reactions were carried out in oven-dried glassware under argon atmosphere. Solvents were purified and dried according to recommended procedures. Thin-layer chromatography (TLC) was carried out on sheets of silica gel 60 F254 on aluminium (layer thickness 0.25 mm, Merck). Visualization of the developed chromatogram was achieved with UV light and ceric ammonium molybdate (CAM) or ninhydrin stains. Chromatographic purification of products was accomplished by column chromatography on silica gels (mesh 100-200 or 230–400). UV/Vis spectra were recorded on Agilent Cary 3500 UV Visible spectrometer. Matrix-Assisted Laser Desorption Ionization (MALDI) mass spectra were recorded on BrukerultrafleXtreme MALDI-TOF/TOF system.

#### Synthesis of Fmoc protected thiourea morpholino active monomer

For the synthesis of GMO (Guanidinium Morpholino Oligonucleotides, GMO) part of GMO-PMO chimera, the Fmoc protected thiourea MMTr-morpholino active monomers of C, A and G nucleosides were required. They were synthesized using the modified protocol as described earlier (8).

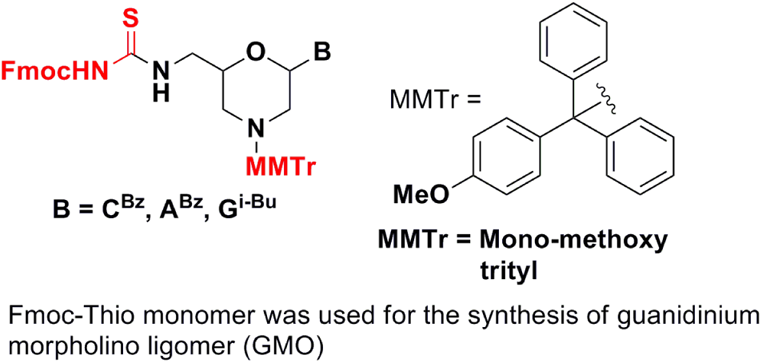

#### Synthesis of Trityl protected chlorophosphoramidate morpholino active monomer

For the synthesis of PMO we have used chlorophosphoramidate monomers of A, T, G and C which was synthesized as per earlier report (9).

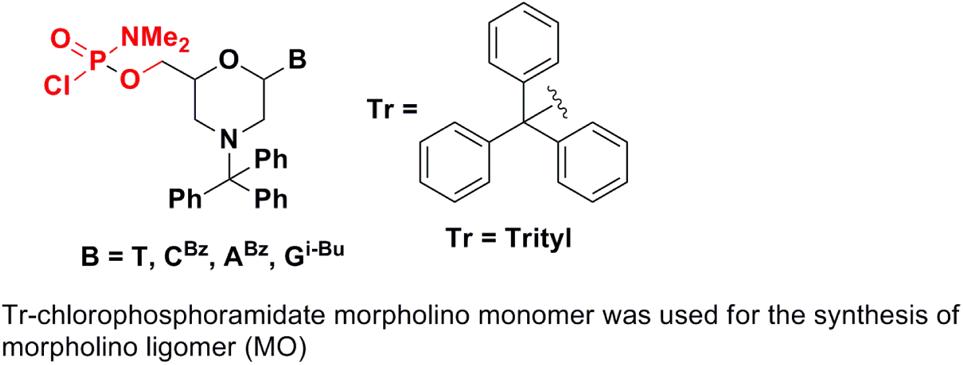

#### Functionalization of solid support with linker and loading monomer

Prior to solid phase synthesis of GMO-PMO sequences, polystyrene solid support was functionalized with aminocaproic acid linker and loading monomer as per earlier report (10).

#### Solid phase synthesis of GMO-PMO (Fig S9 A)

After the successful incorporation of linker and loading monomer on polystyrene solid support, coupling for GMO synthesis was initiated. For GMO synthesis 5 equivalent of Fmoc protected thiourea, 5 equivalents of HgCl_2_ and 5 equivalent of NEM were added in NMP solvent. This step was repeated for another two times in 2 hrs interval. Total coupling time per GMO unit was 6 hrs. Excess reagents were washed with 20 % Thiophenol-NMP and NMP. Unreacted amine was capped with (1:1) mixture of 10 % Ac_2_O-NMP and 10 % DIPEA-NMP. MMTr group was deprotected using the deblocking cocktail (CYPTFA) (10, 11). The synthetic cycle (washing, coupling, capping and deblocking) was repeated for another three GMO monomers (see the structures above) to get the GMO unit. The GMO pentamer was further reacted with chlorophosphoramidate monomer (see the structure above). The morpholino part was synthesized as per our reported method (11). Full length GMO-PMO was cleaved from solid support using 33 % aqueous ammonia at 55°C for 16 hrs and purified by acetone precipitation. Purity of the synthesized GMO-PMOs were checked in HPLC (Fig S9B and C) and characterized by MALDI-TOF. Henceforth GMO-PMO will be denoted as MO.

#### Sequence of AUF1-MO

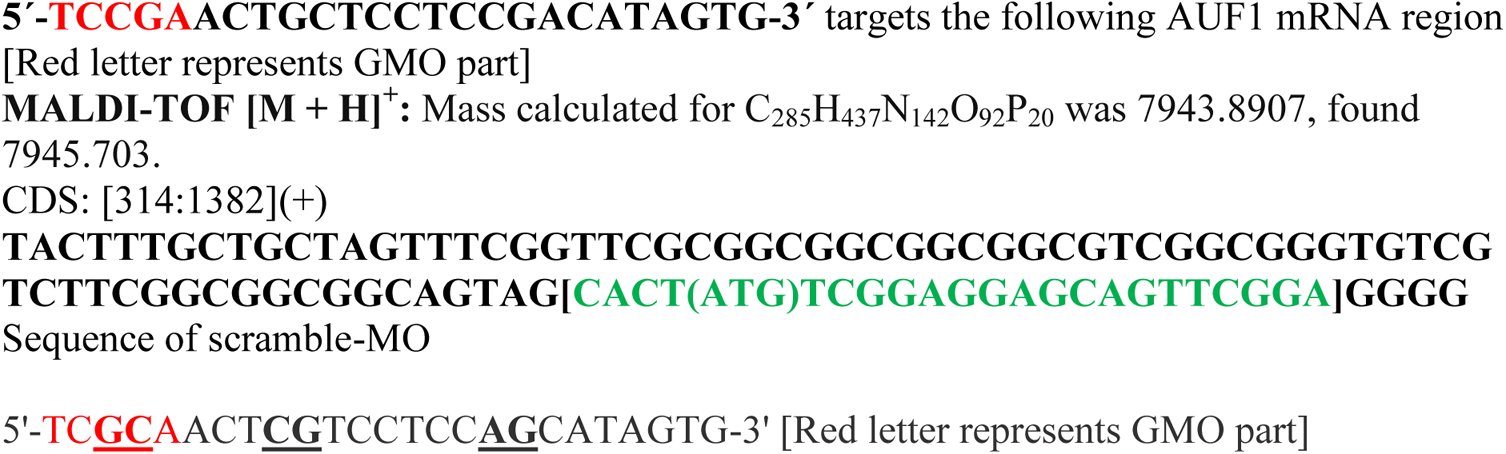

**MALDI-TOF [M + H]^+^:** Mass calculated for C_285_H_437_N_142_O_92_P_20_ was 7943.8907, found 7942.554.

## FIGURE LEGENDS

**Figure S1:**
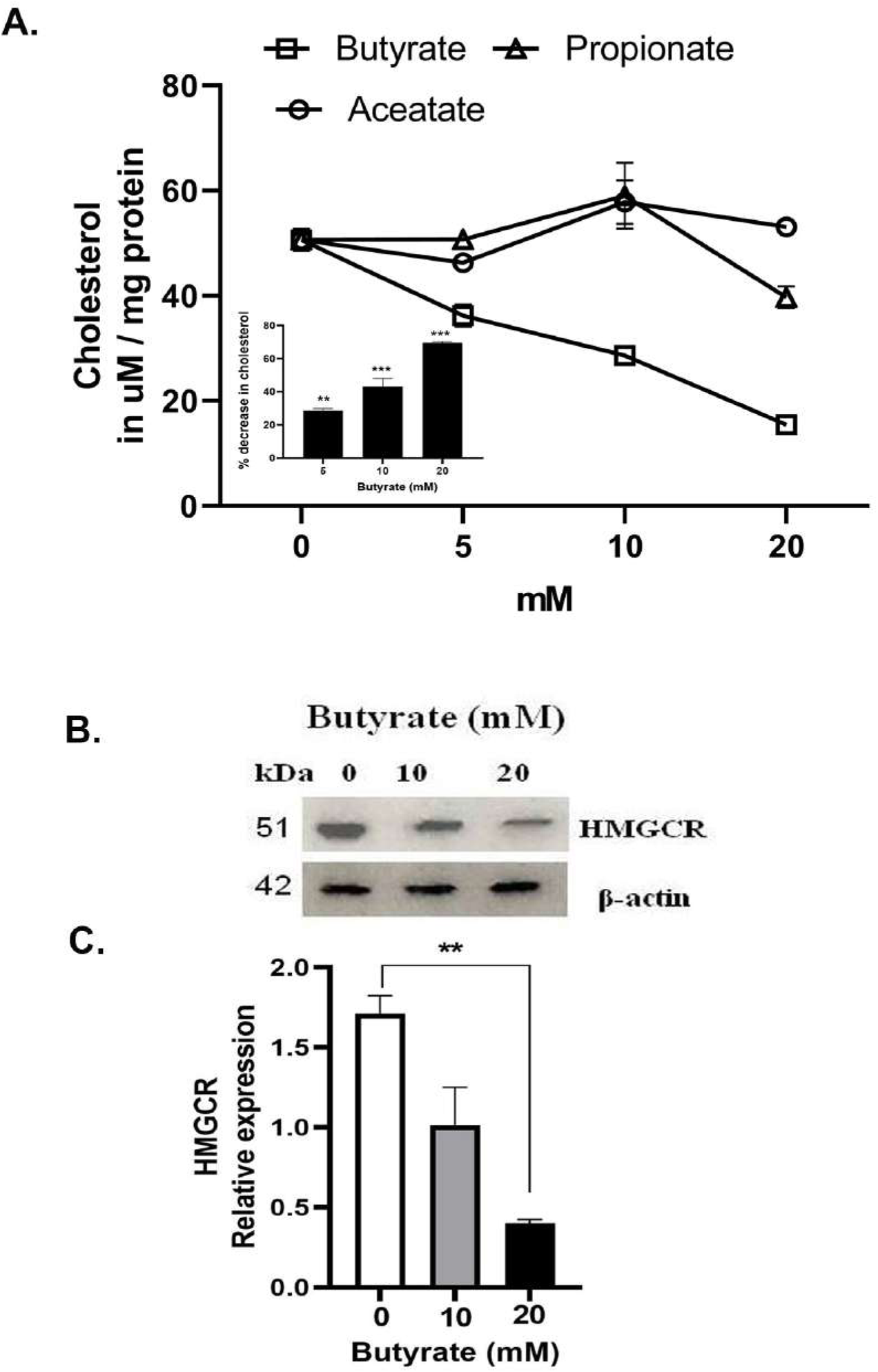
Cellular cholesterol status in Huh7 cells before and after treatment with either butyrate or aceatate or propionate and HMGCR expression with butyrate treatment. Cellular cholesterol status in response to increasing concentration of butyrate or propionate or aceatate (in mM) expressed as µM cholesterol / mg cellular protein (A). Percent decrease in cholesterol with respect to control as a function of butyrate concentration (A, Inset). Western blot of HMGCR expression as a function of butyrate concentration (B). The corresponding densitometry using ImageJ showing relative expression of HMGCR with respect to β-actin control (C). (N=3). *** represents p<0.001 compared to untreated control.

**Figure S2:**
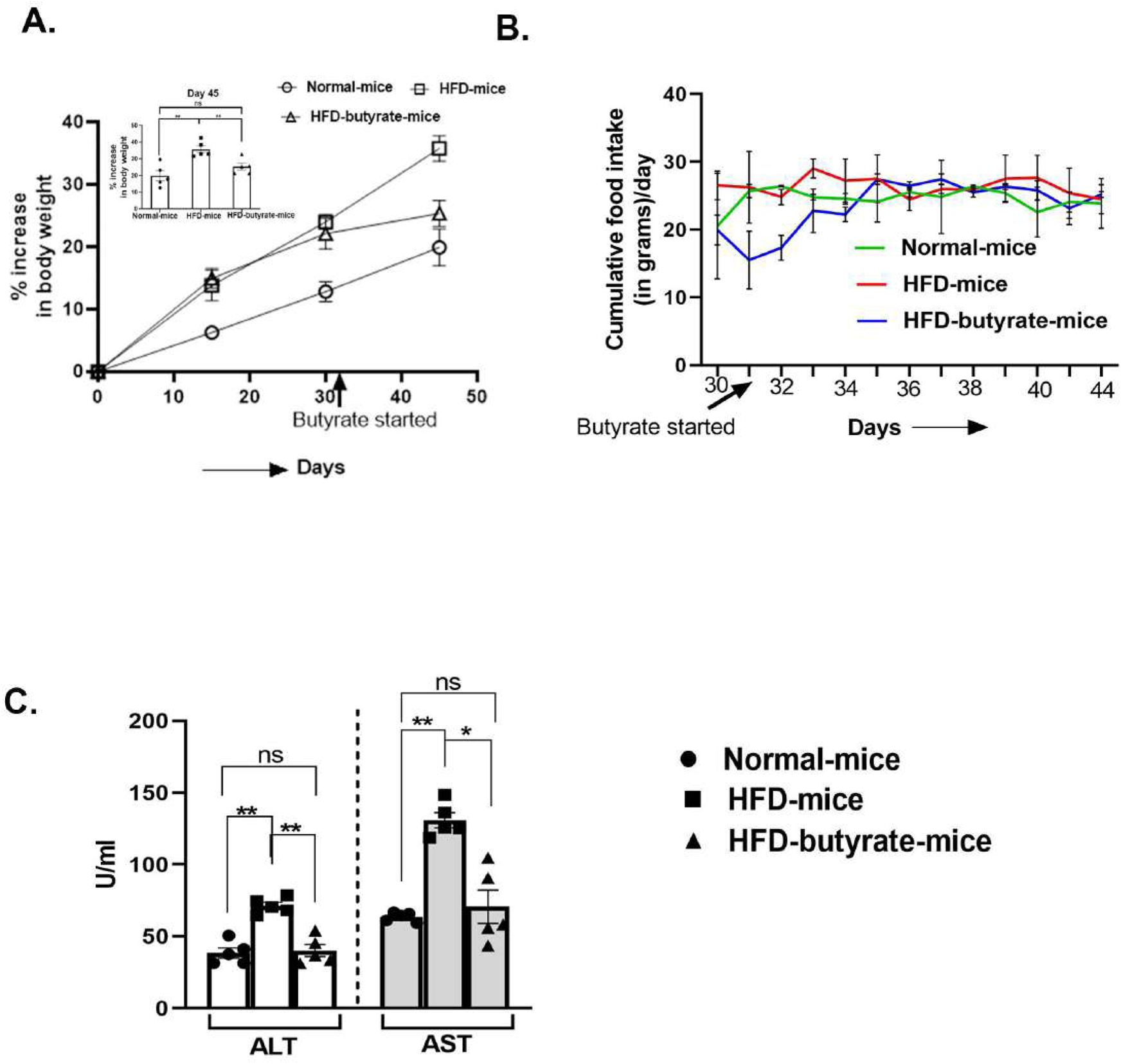
Food consumption, body weight, HDL, LDL status and hepatic enzymes in normal-mice, HFD-mice and HFD-butyrate-mice. Percentage increase in body weight as a function of days (A) and inset showing percent increase in body weight on day 45 (A, inset), cumulative food consumption of 5 mice (in gms) / day was estimated from measuring food every 24 h over a period of 15 days in all the groups. Arrow on the abscissa indicated starting of butyrate treatment. (B), and liver enzymes (ALT & AST) expressed in U/ml (C) were determined in all three groups of mice. N = 5 / group, data is represented as mean ±SE. The experiment was repeated thrice. ** represents p<0.01, * represents p<0.05, ns represents not significant.

**Figure S3:**
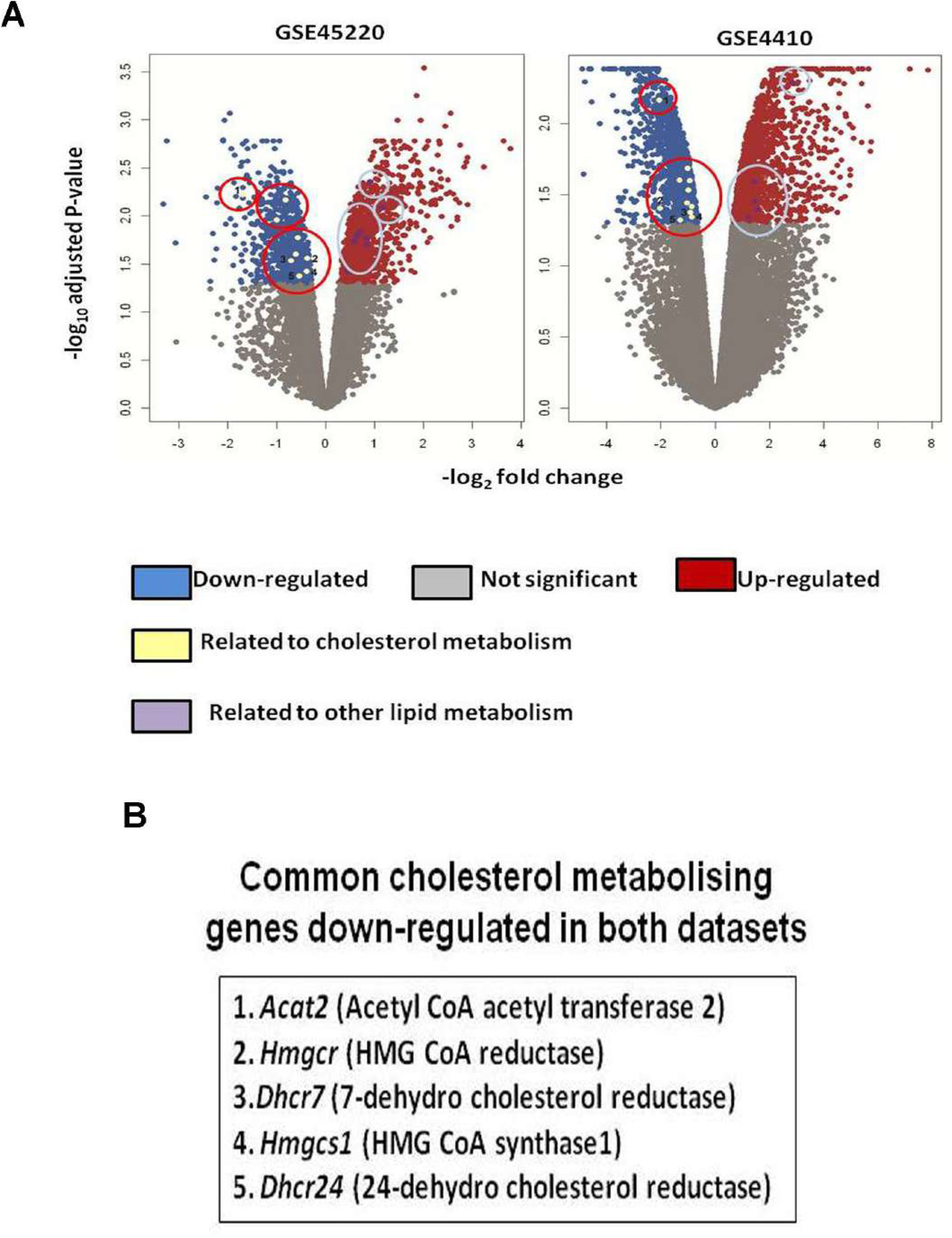
Volcano plot of the publicly available microarray data of control vs butyrate treated HeLa cells (GSE45220) and colon epithelial cell MCE301 (GSE4410) (A). Blue dots, red dots, grey dots represents genes that are down-regulated, up-regulated and non significant respectively. Yellow dots (encircled in red) and purple dots (encircled in light blue) represent cholesterol metabolising and other lipid metabolising genes respectively. The common important genes related to cholesterol metabolism that were down-regulated in both datasets are presented as 1-5 in the box (B).

**Figure S4:**
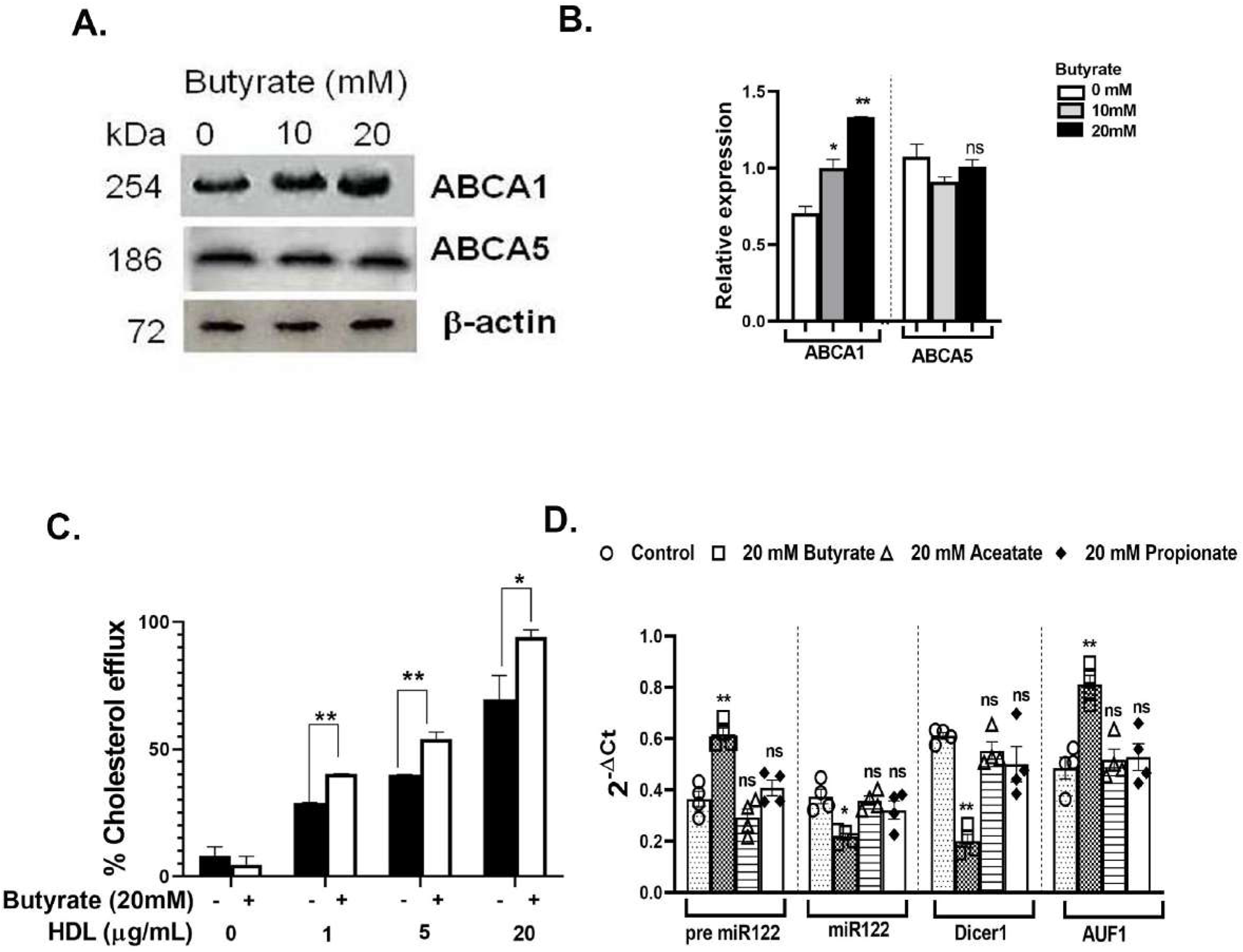
Analysis of ABCA1 and ABCA5 expression in Huh7 by western blot, functional analysis of cholesterol efflux with and without butyrate treatment and status of miR122, Dicer1 and AUF1 in presence and absence of either butyrateor aceatate or propionate. Analysis of expression of ABCA1 and ABCA5 in Huh7 cells by western blot as a function of butyrate concentration (A). The corresponding densitometry using ImageJ showing relative expression of ABCA1 and ABCA5 with respect to β-actin control (B). Percent cholesterol efflux as function of butyrate concentration in the form of 22-NBD-cholesterol was monitored by measuring 22-NBD fluorescence. Huh7 cells were loaded with liposomal 22-NBD-cholesterol for 24 h and subsequently treated with 20mM butyrate. The cells were washed and equilibrated in serum free medium for 18h. Thereafter the cells were treated with or without HDL (1 µg/ml, 5 µg/ml and 20 µg/ml). The fluorescence intensity (FI) of 22-NBD-cholesterol in the medium and cell lysate was detected by MT-600F fluorescence microplate reader (Corona Electric, Hitachinaka, Japan) using 469 nm excitation and 537 nm emission filters in a black polystyrene 96-well plate. The efflux (%) was calculated as (FI_sup_ X 100)/ (FI_sup_ + FI_cell lysate_) (C). Expression of pre-miR122, miR122, Dicer1 and AUF1 (D) after butyrate or propionate or aceatate treatment 20 mM each as measured by qPCR. N ≥3 for all data sets, data is represented as mean ±SE. *** represents p<0.001, ** represents p<0.01, * represents p<0.05, ns represents not significant.

**Figure S5:**
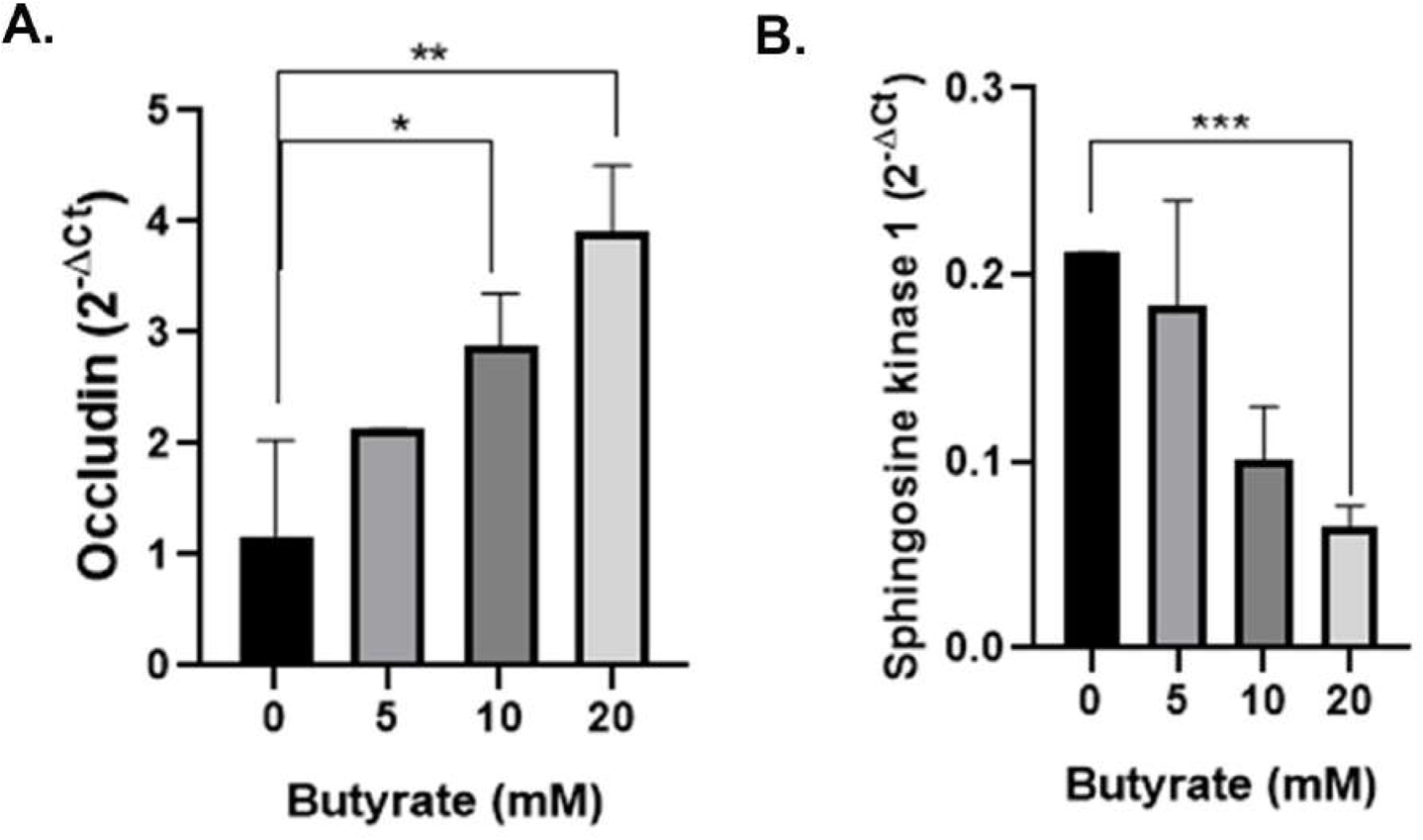
Expression of Occludin (A) and Sphingosine kinase1 (Sphk1) (B) as a function of butyrate concentration in Huh7 cells as measured by qPCR. N=2. The data is represented as mean ± SE. *** represents p<0.001, ** represents p<0.01, * represents p<0.05.

**Figure S6:**
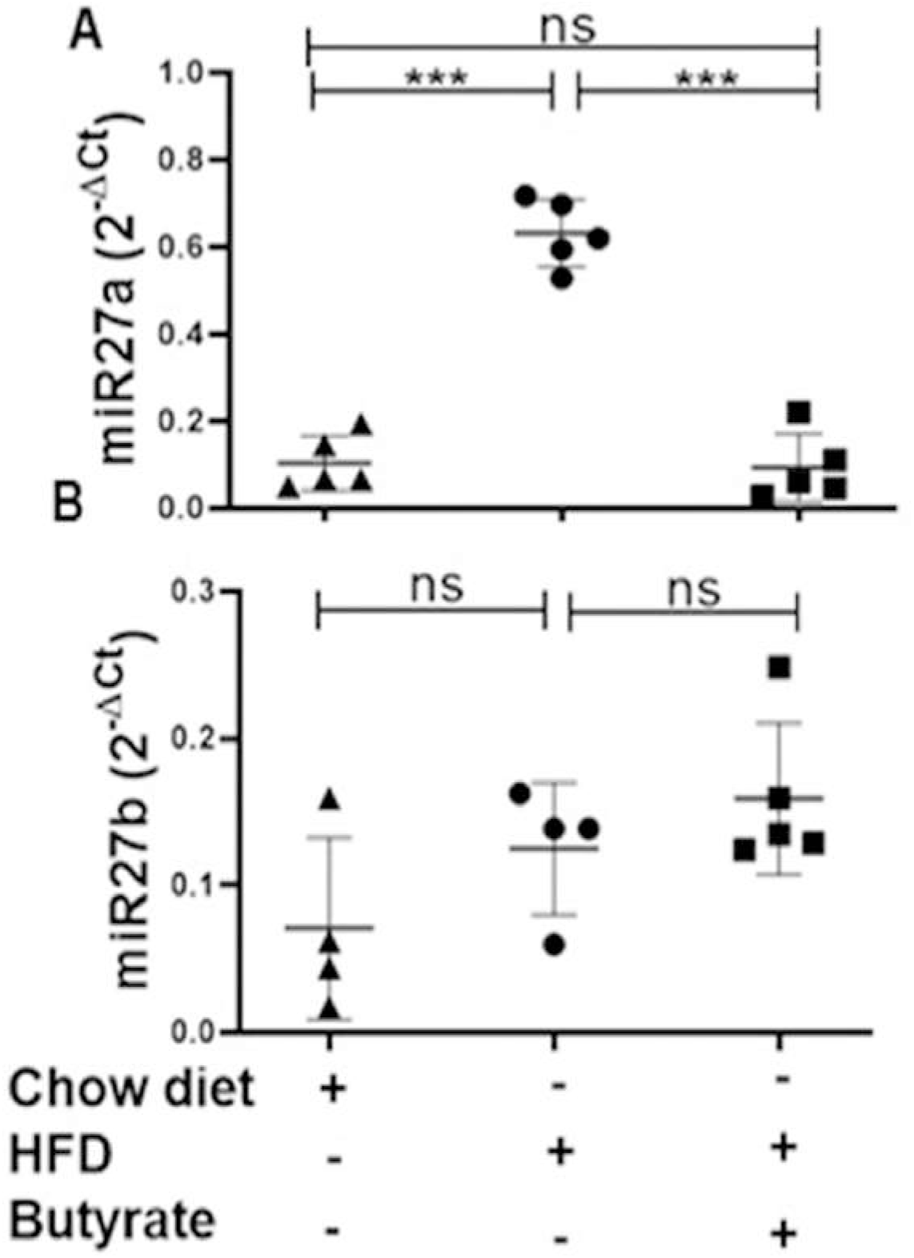
Hepatic expression of miR27a and miR27b in normal-mice, HFD-mice and HFD-butyrate-mice. Hepatic expression of miR27a (A) and miR27b (B) in normal-mice, HFD-mice and HFD-butyrate-mice measured by qPCR. N=4 /group, data is represented as mean ±SE. *** represents p<0.001, **represents p<0.01, * represents p<0.05.

**Figure S7:**
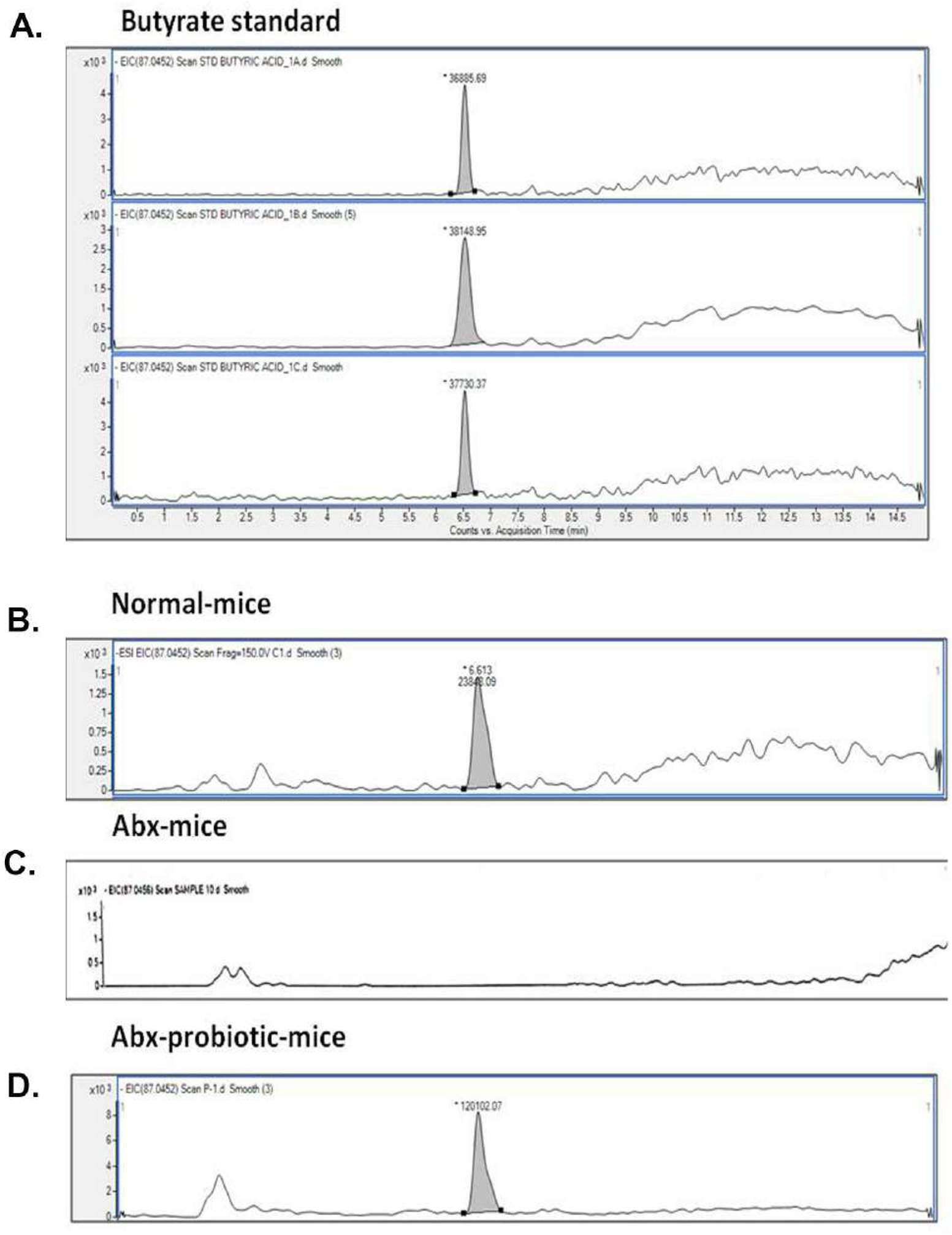
Determination of fecal butyrate by LC-MS. Representative peak of butyrate in LC-MS. Standard (2 mg/ml) (A), Normal-mice (B), Abx-mice (C) and Abx-probiotic-mice (C).

**Figure S8:**
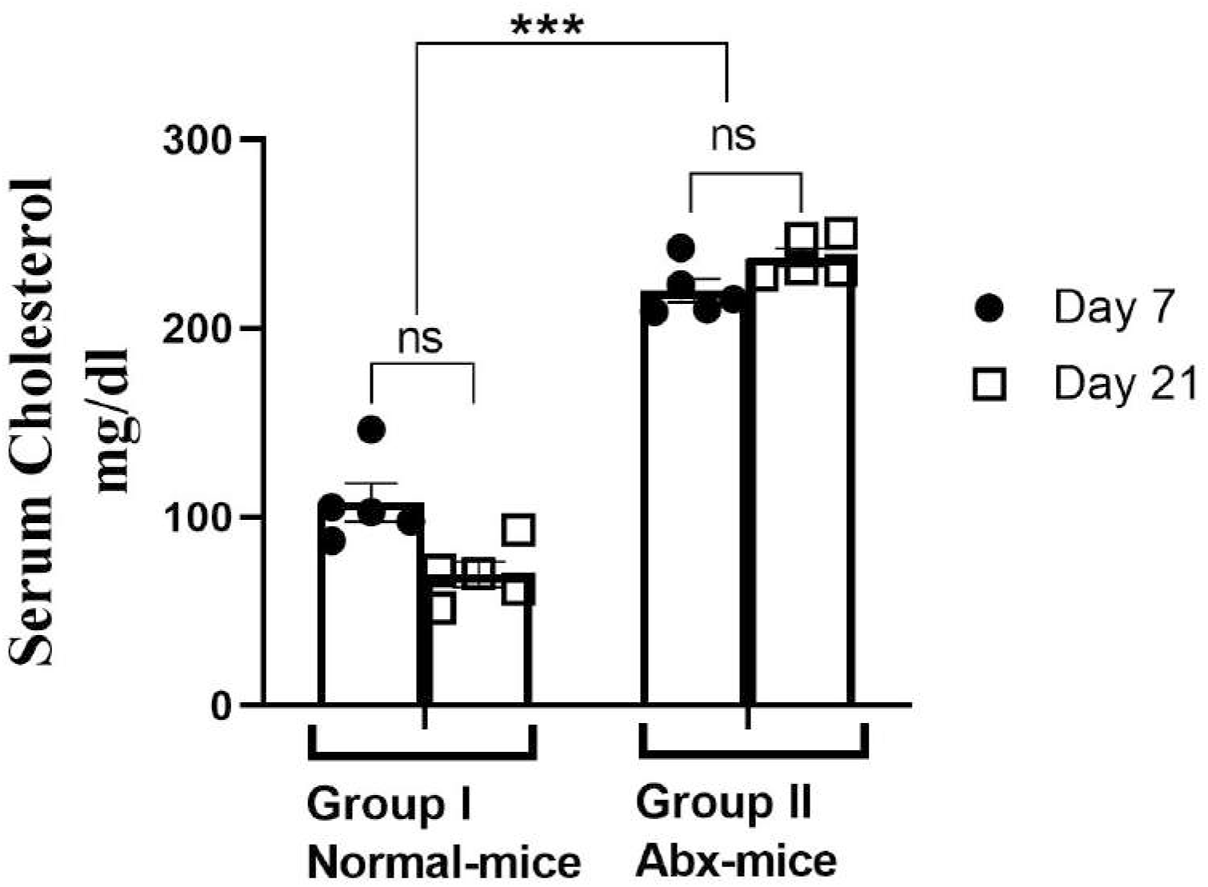
Antibiotic treatment increases serum cholesterol. The serum cholesterol of normal-mice (A) and Abx-mice (B) on day 7 and day 21.

**Figure S9:**
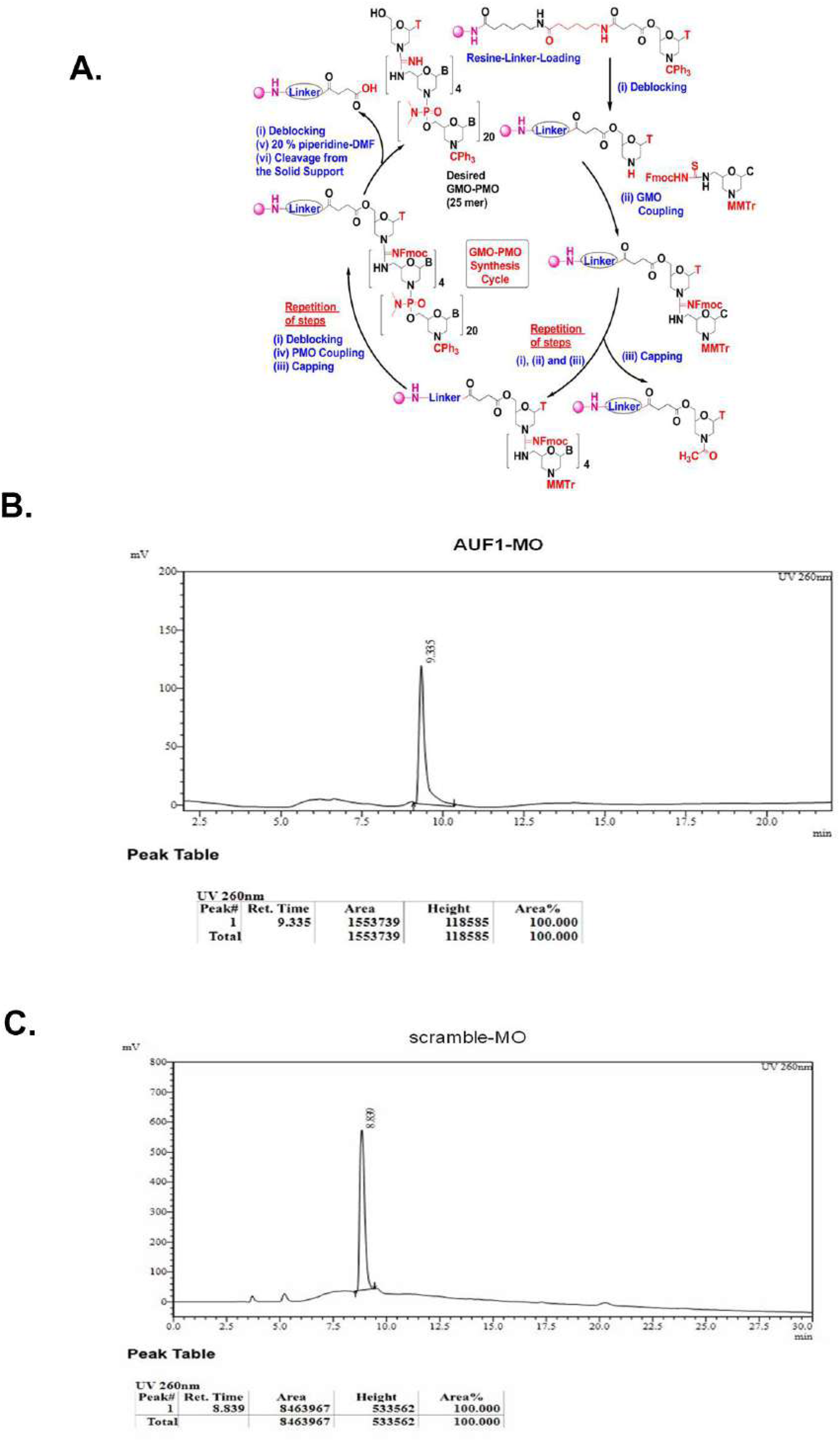
Synthesis cycle of GMO-PMO (MO) and characterization of scramble-MO and AUF1-MO by HPLC. Synthesis cycle of GMO-PMO (MO) (A), HPLC purity peak for AUF1-MO (B) and scramble-MO (C).

**Figure S10:**
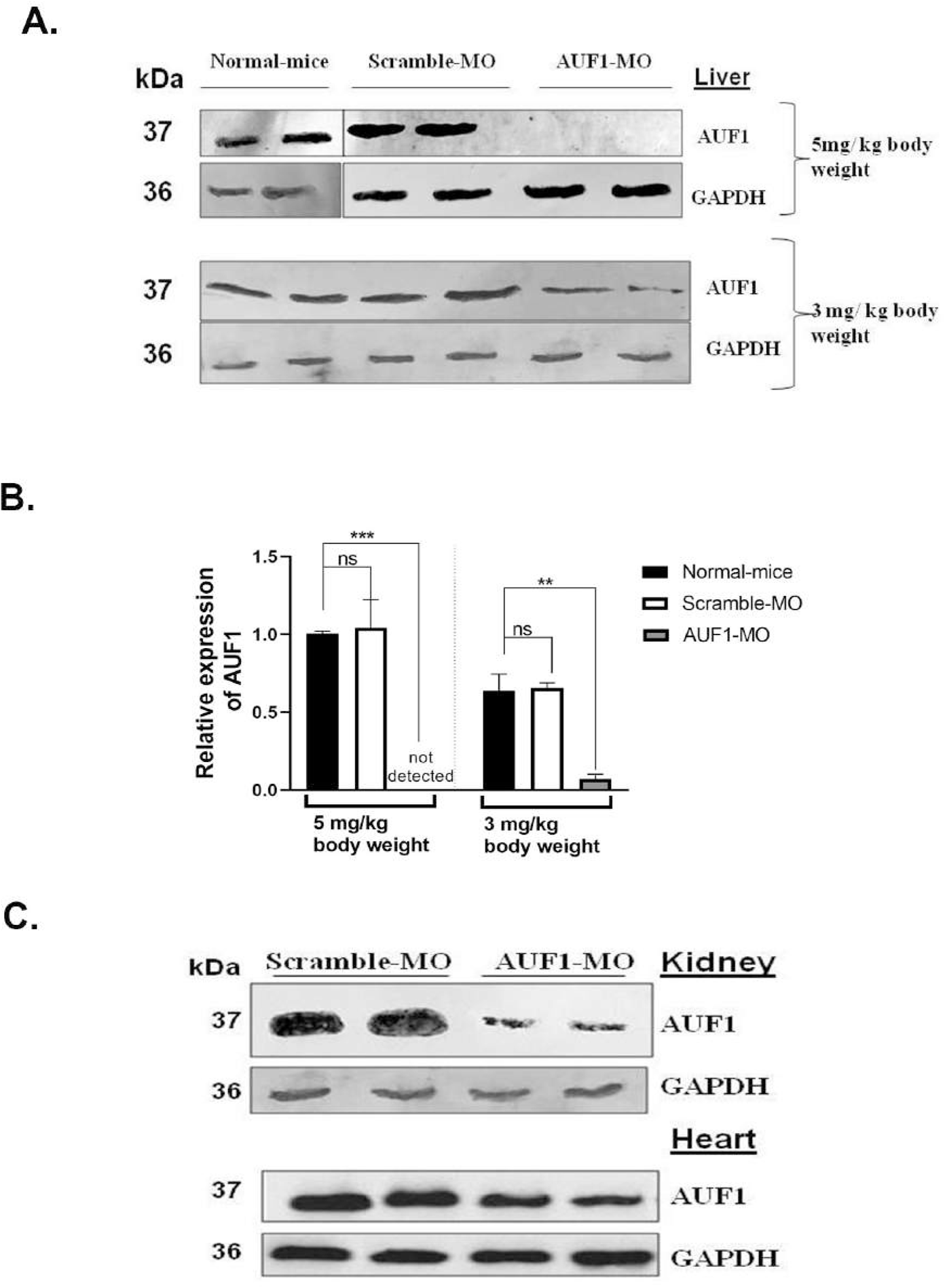
Specificity of AUF1-MO in selective knockdown of AUF1 in mice. Mice were injected in the tail vein with either 5 mg/kg body weight or 3 mg/kg body weight of AUF1-MO/ scramble-MO. The liver samples were collected 7 days post injection and western blot analysis was performed to determine the expression of AUF1 (A) and corresponding densitometric analysis of AUF1 expression by ImageJ (B). Western blot of AUF1 in heart and kidney of mice receiving AUF1-MO (C). GAPDH was used as control for all experiment. The anti-AUF1 antibody (Cell signalling) used in this study was specific to AUF-1^p37^. N=2

